# Dynamic interaction between tentacle-tethered microdomains in Integrator and NELF regulates the promoter-proximal pause checkpoint

**DOI:** 10.64898/2026.06.30.735463

**Authors:** Daniel Blears, Pernille Bech, David Lopez Martinez, Pengfei Dai, Riyaz A. Shah, Nova Fong, Zhong Han, Richard Mitter, Ryan Sheridan, Jens Vilstrup Johansen, A. Barbara Dirac-Svejstrup, David Bentley, Jesper Q. Svejstrup

## Abstract

Negative Elongation Factor (NELF) and Integrator are general regulators of transcriptional pausing and premature termination at the beginning of protein-coding genes. While NELF is a key pausing factor, Integrator triggers premature termination of dysfunctional RNAPII complexes. Here, we identify a direct interaction between the plant homeodomain (PHD) finger of Integrator subunit 12 (INTS12) and a previously unrecognized C-terminal domain (CTD) in the flexible tentacle of NELF-A. Mutation of either domain disrupts the interaction, and although RNAPII-associated pausing complexes still form, they are dysfunctional. Importantly, while deletion of the INTS12 PHD finger functionally mimics Integrator loss, a single point mutation in NELF-A CTD instead mimics loss of NELF, suggesting the presence of competitive interactions. Together, our findings uncover a novel mode of PHD finger binding and outline how an interaction between Integrator and NELF regulates transcriptional pausing and premature termination.

## Introduction

In higher eukaryotes, transcription of protein-coding genes involves a fundamental and critical pausing step shortly downstream of the transcription start site (TSS), known as promoter-proximal pausing^1,2^. After pausing, RNA polymerase II (RNAPII) complexes then either progress into productive transcript elongation or undergo premature termination^3–6^. DRB-sensitivity inducing factor (DSIF), Negative Elongation Factor (NELF) and Integrator have important functions in regulating pausing and premature termination in promoter-proximal regions^7–13,1,14,2,15^, yet the mechanisms governing their function remain unclear. In the predominant model, NELF, DSIF and RNAPII form a stable pausing complex, while Integrator associates dynamically with this stable complex to promote premature transcription termination^1^.

Structural and biochemical analysis of paused RNAPII bound by Integrator, NELF and DSIF has partially revealed how these complexes assemble and interact^16,12,17^. However, while cryo-electron microscopy (cryo-EM) approaches have charted the majority of the protein sequences^16,12,17,18^, certain peripheral regions have not been visualized, partly due to being structurally dynamic or disordered. For example, structural and functional information regarding the flexible “tentacle” domains of the NELF-A and -E subunits, as well as the entire Integrator complex subunit 12 (INTS12), is missing.

According to *in silico* structural predictions, the INTS12 subunit of Integrator is mostly disordered, yet harbors a domain belonging to the Plant Homeodomain (PHD) finger family; hence its alternative name: PHD finger protein 22 (PHF22). The PHD family encompasses conserved domains with diverse functions in chromatin regulation, and most characterized family members interact directly with histones, particularly the regulatory N-terminus of histone 3 (H3)^19–24^. PHD domains have thus gained a reputation as a family of epigenetic readers, with their histone binding capabilities allowing the tethering of protein complexes to specifically modified nucleosomes, for example to achieve gene regulatory outputs^23^.

This well-studied function of PHD finger domains prompted our investigation into the function of INTS12. Here, we show that INTS12, and specifically its PHD finger domain, are critical for correct processing of RNAPII at the promoter-proximal pause and for the formation of competent elongation complexes. Remarkably, the INTS12 PHD finger domain is required - not for binding histones - but for interaction with a previously uncharacterized NELF microdomain, located in the distal, C-terminal extension of the flexible NELF-A tentacle. This reveals an interesting type of interaction between two small domains, each tethered to their respective core complexes by flexible linkers. We demonstrate the competitive nature of this interaction with transcriptional activators and propose that the slow association of these tethered satellite microdomains may represent a kinetic quality-control step at the promoter-proximal pause. Together, our findings suggest a novel mechanism of transcription regulation by Integrator and NELF, which expands our conception of the function of the PHD finger domain and reconciles the paradoxical functions of Integrator as both an attenuator and activator of transcription.

## Results

### INTS12 is a key Integrator component, but not required for complex assembly

Given that INTS12 contains a PHD finger domain, we postulated that it might facilitate recruitment of Integrator to pausing complexes in chromatin, perhaps at least partly through interaction with nucleosomes. We therefore began by assessing the effect of siRNA-mediated INTS12 depletion on Integrator levels in chromatin from HEK293 Flp-In TREx cells (Figure 1A). Interestingly, INTS12 loss had no obvious effect on the levels of various other Integrator subunits in chromatin, whereas similar knockdown of INTS4 or INTS6 had strong effects as described previously^25^ (Figure 1A, compare lane 8 with lanes 5-7). In addition, we analyzed the Integrator interactome via INTS11 immunoprecipitation (IP) and mass spectrometry, in the presence or absence of INTS12. The loss of INTS12 consistently had little or no effect on the integrity of the Integrator complex or its association with chromatin and RNAPII (Figure 1B), suggesting that INTS12 is neither required for assembly of the core Integrator complex nor its recruitment to paused polymerase complexes. These findings are consistent with cryo-EM data in which the INTS12 subunit has remained unresolved^12,17,18^, suggesting it is a peripheral subunit with high mobility relative to the Integrator core.

**Figure 1.**
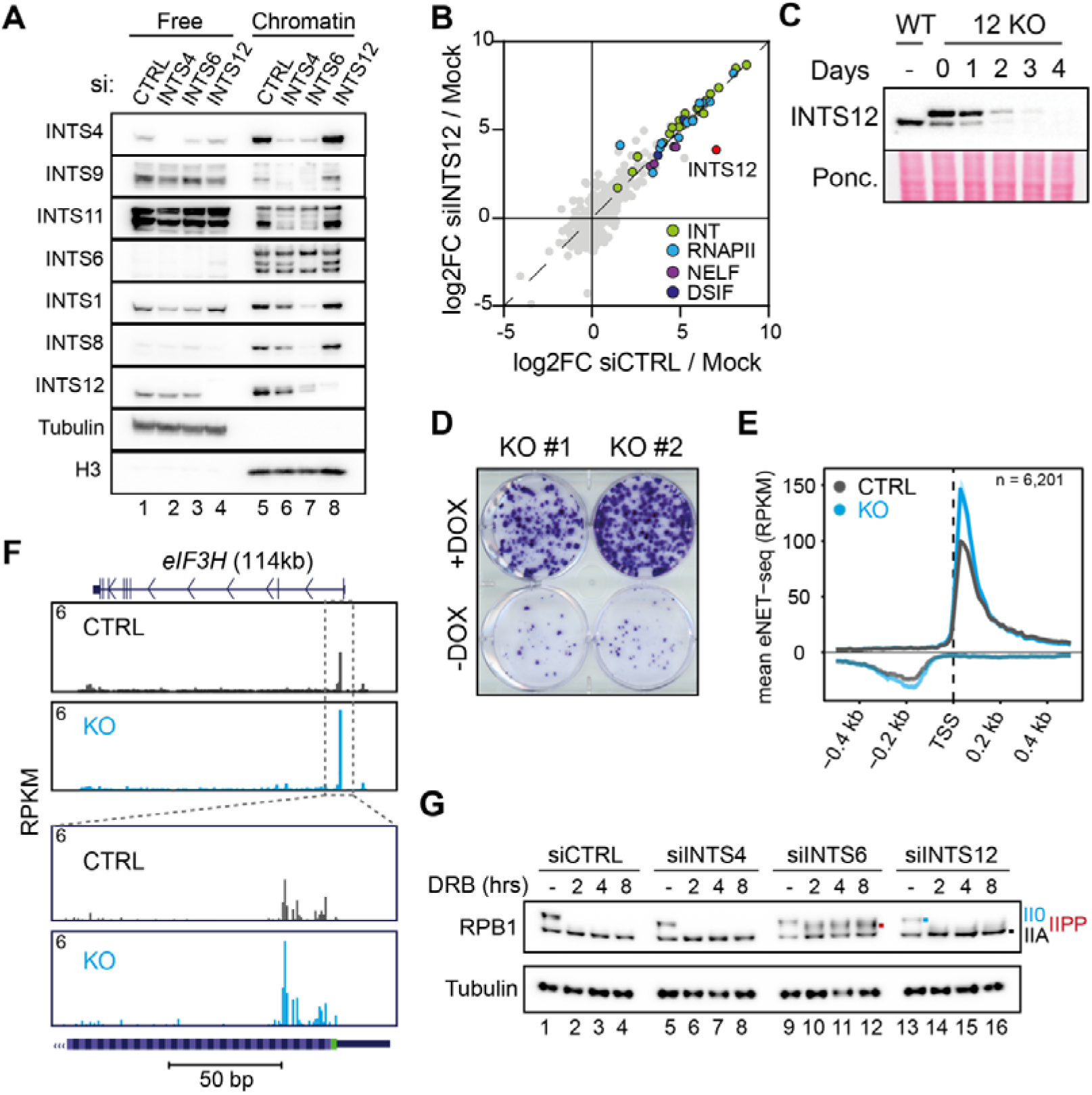
Effect of INTS12 on promoter-proximal pausing. **A.** Western blot analysis of Integrator composition in chromatin (lanes 5-8) upon siRNA depletion of the indicated factors. ‘Free’ indicates protein not associated with chromatin. **B.** The INTS11 interactome in WT cells expressing FLAG/HA-tagged INTS11, compared to the same cells after siRNA kd of INTS12. IPs are shown as log2FC compared with negative “mock” control (parental cells without FLAG-INTS11 expression). **C.** Western blot analysis with INTS12 antibody of whole cell extracts from WT cells and *INTS12* KO rescue cells expressing FLAG/HA-tagged INTS12 from a DOX-regulated promoter after removal of DOX from the culture medium. Note that, in addition to the main isoform, a minor INTS12 isoform is also observed in the presence of DOX, with a mass similar to that of endogenous INTS12. This form (possibly INTS12 from which the tag has been proteolyzed) disappears after DOX removal, showing that it is derived from FLAG/HA-tagged INTS11 and not an endogenous gene product. **D.** Single-cell survival assay of *INTS12* KO cells with or without DOX. **E.** Average eNET-seq profiles for protein-coding genes in WT (CTRL, Dox present) and *INTS12* KO (DOX removed), centered on the transcription start site (TSS). **F.** Single gene example of data in E. **G.** Western blot analysis of RPB1 upon siRNA depletion of the indicated factors, with the different RPB1 isoforms indicated on the right and with matched colored dots inside the blot. See also Supplementary Figure S1.

Nevertheless, we were unable to obtain stable *INTS12* knock-out (KO) clones by CRISPR in HEK293 cells (data not shown), providing a first suggestion that the *INTS12* gene is essential for cell fitness and survival. Instead, we first integrated a doxycycline (DOX)-inducible transgene expressing a FLAG/HA-tagged version of INTS12 into cells and subsequently knocked out the endogenous *INTS12* gene by CRISPR, while maintaining INTS12 protein levels from the transgene in the presence of DOX. Using this approach, we now obtained cells lacking the endogenous *INTS12* gene, which in the presence of DOX expressed the tagged INTS12 version at levels similar to those of wild type cells. Using this system, we were able to deplete INTS12 by removing DOX from the cell culture medium, effectively generating an INTS12 knock-out on demand (Figure 1C). Importantly, after depletion of INTS12, cells lost their ability to form clonal colonies (Figure 1D), showing that INTS12 is indeed required for optimal cell fitness and survival in HEK293 cells.

We then investigated the effects of INTS12 depletion on transcription using (enhanced) mammalian nascent elongating transcript sequencing (eNET-seq)^26^ to map the occupancy of RNAPII complexes genome-wide. eNET-seq metagene analysis revealed sharp peaks of RNAPII density at the 5’ ends of protein-coding genes (Figure 1E, F): a signature attributed to promoter-proximal pausing^26,25^. INTS12 loss resulted in a clear increase in promoter-proximal RNAPII density (Figure 1E), while the density in gene-bodies and 3’ ends was slightly reduced (Supplementary Figure S1A). Increased pausing was also quantified by calculating the fraction of paused reads, otherwise known as “pausing propensity”^26^. In *INTS12* KO cells, this fraction was specifically increased immediately downstream of the transcriptional start site (TSS) but not generally in gene bodies (Supplementary Figure S1B). Loss of INTS12 thus results in stabilization of RNAPII in the promoter-proximal region, a well characterized phenotype of cells with defective Integrator^27,28^.

We recently showed that upon treatment with 5,6-dichloro-1-beta-D-ribofuranosylbenzimidazole (DRB), RNAPII is dephosphorylated in an INTS6-dependent manner, resulting in marked loss of hyperphosphorylated RNAPII (IIO) and concomitant accumulation of the unphosphorylated (IIA) form^25^. Removal of INTS6, and consequently Integrator phosphatase activity, prevents dephosphorylation of RNAPII and instead stabilizes a paused, serine-5 phosphorylated intermediate (named IIPP)^25^. We assessed RNAPII levels upon DRB treatment in INTS12 depleted cells and did not observe stabilization of IIPP, suggesting that the Integrator phosphatase activity is functional in the absence of INTS12 (Figure 1G). Instead, an accumulation of unphosphorylated RNAPII (IIA) relative to control cells was observed after DRB treatment (Figure 1G, compare lanes 14-16 with 2-4). This is consistent with the idea that turnover of RNAPII at the promoter-proximal pause is decreased in these cells.

### INTS12 loss results in transcript elongation defects

We next carried out nascent transcription profiling using transient-transcriptome sequencing with chemical RNA fragmentation (TT_chem_-seq)^29^ for assessing transcription patterns after INTS12 loss. This showed that INTS12 loss results in a global reprogramming of transcription, with relatively more transcription in the 5’ ends of genes and less activity at their 3’ ends (Figure 2A-C), indicative of a defect in transcript elongation. Similar results were obtained after siRNA-mediated knockdown of INTS12 (Supplementary Figure S1C). Calculation of the log2 fold-change difference between INTS12 knockdown and control samples showed a sharp reduction in the relative transcript abundance as RNAPII moves away from the TSS and into gene bodies (Figure 2C), again in agreement with a defect in elongation speed and/or processivity. Together, these results suggest that without INTS12, RNAPII dwells longer at the promoter-proximal pause and elongation complexes are released into the gene body that are somehow transcriptionally compromised. Interestingly, when we looked at a subset of short genes (<14kb), the average density plots showed a relative increase in transcription across the full length of these genes (Figure 2D and Supplementary Figure S1D). This could suggest that some short genes escape transcript elongation defects that accumulate with distance, and in fact become upregulated. This is consistent with the phenotype previously reported after loss of Integrator RNA cleavage activity^30,27^. Other recent results show that Integrator deficiency also results in upregulation of certain short genes in the integrated stress response^31^.

**Figure 2.**
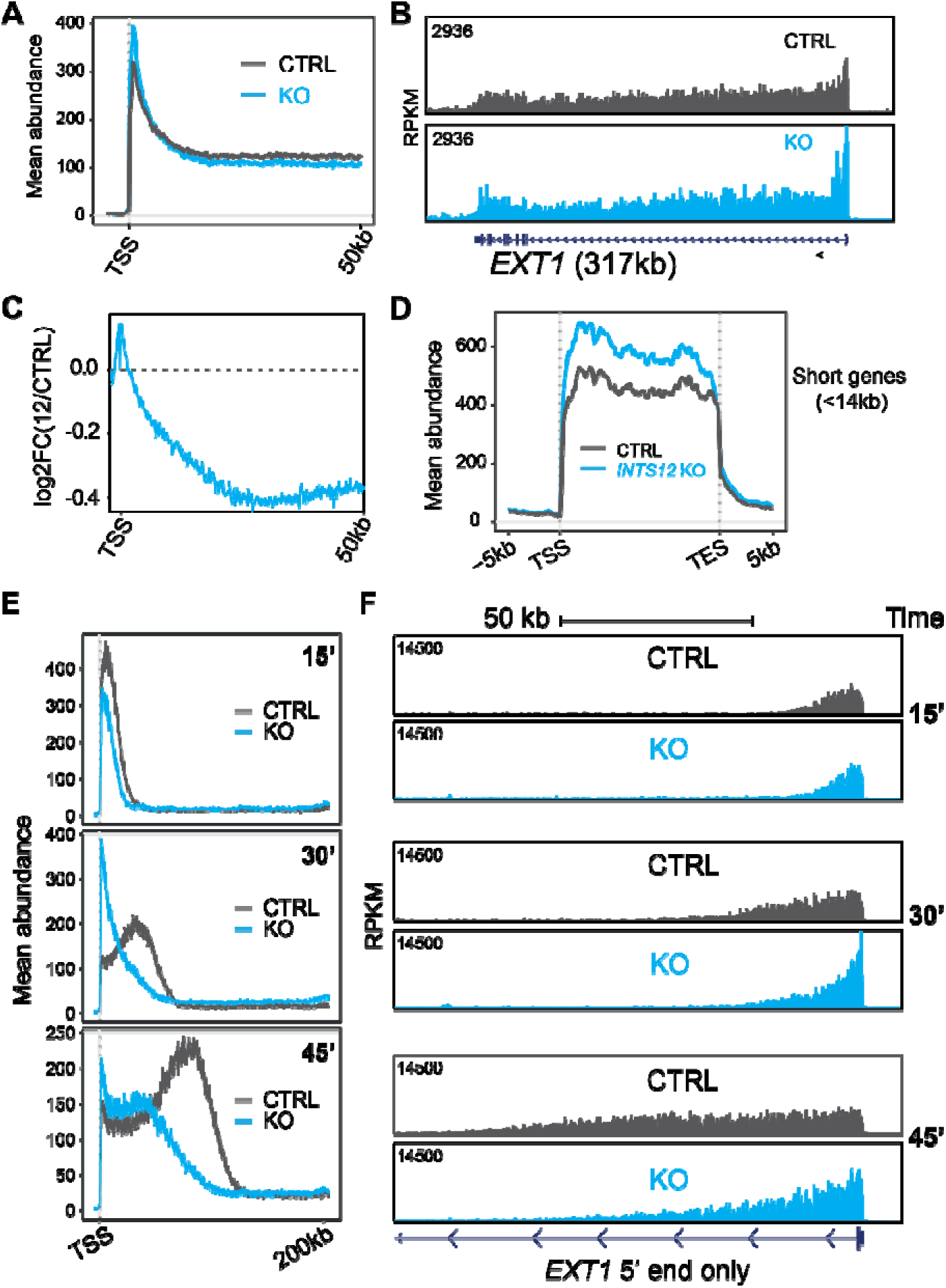
The effect of INTS12 on transcription. **A.** Average TTchem-seq profiles for protein-coding genes >50kb, centered on the TSS and extending to 50kb into gene bodies. **B.** TTchem-seq profile for the long *EXT1* gene. **C.** As in A, but showing the log fold-change comparison of CTRL and *INTS12* KO cells. **D.** Average full length TTchem-seq profiles for protein-coding genes <14kb. **E.** DRB/TT_chem_-seq: average TT_chem_-seq profiles for protein-coding genes >200kb after DRB washout, centered on the TSS and extending 200kb into gene bodies. Respective time-points post-DRB washout are indicated in bold. **H.** Single gene example of *EXT1* in both CTRL and *INTS12* KO cells. See also Supplementary Figure S1.

To further investigate transcript elongation characteristics in the absence of INTS12, we measured elongation employing a DRB/TT_chem_-seq approach^29^. In this approach, pause-release is inhibited by the addition of DRB for 2 hours, thereby blocking RNAPII progression beyond promoter-proximal regions, while allowing already gene-associated polymerases to run off. DRB is then washed out, allowing synchronized release of RNAPII into transcript elongation, and transcription activity is measured by TT_chem_-seq at 15’, 30’ and 45’ time-points.

By focusing on long genes in which the polymerase cannot reach the gene end (TES) over the course of the experiment, we observed a clear progression of RNAPII into gene bodies after DRB washout in WT cells (Figure 2E, F). However, after loss of INTS12 there was a striking defect in RNAPII progression, with most activity restricted to the 5’-end of genes, even at the 30’ and 45’ time-points (Figure 2E, F). This indicates dramatically slowed elongation in the absence of INTS12, likely due to release of incompetent elongation complexes.

In addition to their function at the promoter-proximal pause, some Integrator subunits are required for the 3’ processing of small-nuclear RNA (snRNA) transcripts and subsequent transcription termination, while others are not^32,33,12,28,25^. Loss of INTS12 had little or no effect on snRNA transcription termination by TT_chem_-seq, while knockdown of the RNA cleavage-module subunit INTS4 resulted in a strong snRNA read-through effect as expected (Supplementary Figure S1E, F). snRNA read-through was also measured using a GFP reporter system, where GFP expression indicates transcriptional read-through resulting from Integrator dysfunction^33^. Again, knockdown of INTS12 had only a marginal effect (Supplementary Figure S1G). Thus, INTS12 function appears to be specific to protein-coding gene promoters and is not essential for snRNA processing.

### The PHD finger domain is required for INTS12 / Integrator function

With the finding that the INTS12 subunit is critical for the Integrator function at protein-coding genes, we aimed to determine the importance of the PHD finger domain. According to Alphafold structure predictions, INTS12 in isolation is mostly disordered, except for a small region at the N-terminus and the PHD finger domain. Structure predictions of the PHD finger show an architecture similar to that of many canonical PHD fingers (Supplementary Figure S2A-D). Characteristic features include two short anti-parallel β-sheets, which in other proteins typically interact with the histone H3 tail to form a triple β-sheet structure (Supplementary Figure S2B) and two Zn^2+^ co-ordination centers formed by cysteine and histidine residues that are highly conserved across all PHD finger domains^23^. We noticed that the INTS12 PHD contains a unique extended N-terminus harboring an α-helix (Supplementary Figure S2A, S2B, S2E); a feature which is somewhat unconventional of PHD fingers and might indicate an extended surface capable of forming an interaction that differs from typical H3 tail binding. Furthermore, analysis of the sequence and structure of the INTS12 PHD finger effectively rules out the possibility of it binding to trimethylated H3K4 (H3K4Me3), as it lacks the canonical “aromatic cage”, a hydrophobic pocket formed by one or more aromatic residues, characteristic of all known H3K4Me binders^23^. Rather, the INTS12 PHD finger more closely aligns with PHDs that are known to bind unmethylated H3K4 (H3K4Me0), in that it harbors a leucine residue at position 1 of the first β-strand; a signature which is conserved among the sub-type of H3K4Me0 binders (Supplementary Figure S2E).

To investigate the functional importance of the INTS12 PHD domain, we constructed cell lines with inducible expression of siRNA-resistant INTS12 (WT) or INTS12 in which the PHD finger domain was deleted (Supplementary Figure S3A). Using this system, we knocked down endogenous INTS12 by siRNA and re-introduced either WT INTS12 or INTS12 lacking the PHD finger (ΔPHD-INTS12) (Supplementary Figure S3B). TT_chem_-seq analysis showed that ΔPHD-INTS12 expression resulted in transcription defects that are similar to complete INTS12 loss, with more RNAPII transcription activity near the promoter-proximal pause, but less activity further into the coding region (Figure 3A, right; and Supplementary Figure S3C). The similarity was particularly obvious in the difference maps (Figure 3A, lower). These data highlight the importance of the PHD finger for INTS12 function.

**Figure 3.**
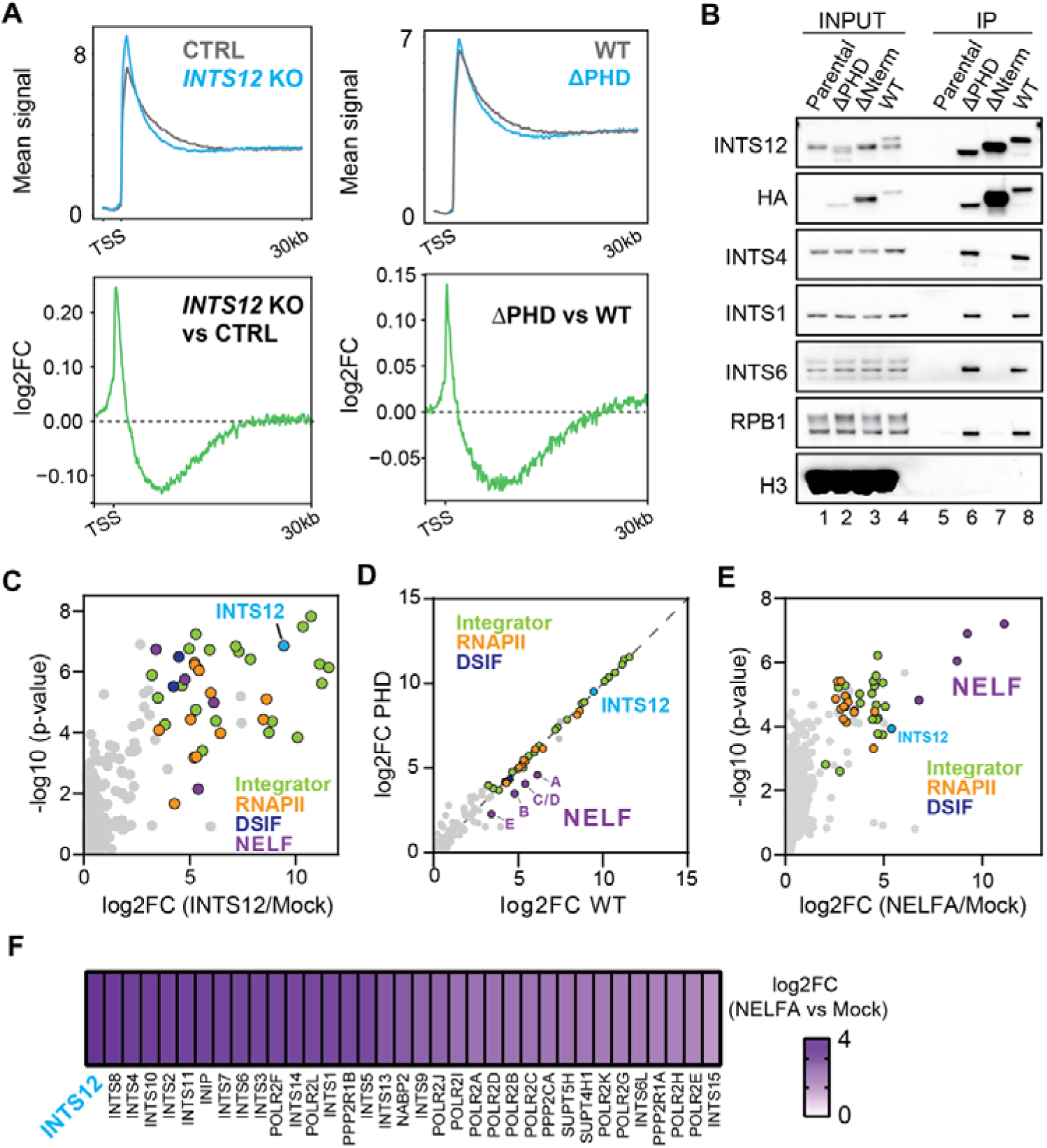
The effect of PHD finger deletion. **A. Upper,** Average TTchem-seq profiles for protein-coding genes >30kb, centered on the TSS and extending to 30kb into gene bodies. *Left,* siRNA mediated INTS12 loss versus control siRNA (CTRL). *Right,* INTS12 ΔPHD vs INTS12 WT in the switchover system. **Lower,** the log fold-change comparisons of the plots above. **B.** Western blot of IP experiments where FLAG/HA-tagged INTS12 WT, ΔPHD, and ΔNterm mutants were pulled down from solubilized chromatin extracts. Histone 3 (H3) was overexposed to show the absence of H3 in the pull downs. **C.** Proteomic analysis of FLAG-tagged INTS12 in chromatin. Log fold-change of pull down and negative non-FLAG control (Mock) is shown, along with the statistical significance (-log10(p-value)). **D.** Comparison of INTS12 WT and ΔPHD pull-downs (shown as log2 fold-change relative to negative control pull down from non-FLAG parental cells). NELF, RNAPII and Integrator subunits are indicated. A line is plotted through x=y for reference. **E.** Proteomic analysis of FLAG-NELFA WT in chromatin. Log fold-change of pull down and negative non-FLAG control (Mock) is shown, along with the statistical significance (-log10(p-value)). **F.** Heat-map showing the log2FC of selected NELFA interactions, focused on subunits of Integrator and the RNAPII elongation complex. See also Supplementary Figures S2 and S3.

We now investigated how loss of the PHD finger affects the interactome of INTS12. WT, ΔPHD-INTS12, or INTS12 lacking the N-terminal 50 amino acids (ΔNterm) (See Supplementary Figure S3A and S3B) were immunoprecipitated from nuclease-solubilized chromatin extracts and the eluates analyzed by Western blotting (Figure 3B) and mass spectrometry (Figure 3C and D; see also Supplementary Figure S3D). As expected, an abundance of Integrator and RNAPII, as well as both the NELF and DSIF were detected in the IPs with WT INTS12 (Figure 3B and C). Intriguingly, however, we were consistently unable to detect significant levels of histones in the INTS12 pulldowns. While it is impossible to make strong conclusions based on this negative result, the apparent lack of histone interaction is consistent with a previous high-throughput study in which the binding of different PHD domains to a series of histone substrates failed to identify specific INTS12-histone tail interactions^24^.

The N-terminal domain of INTS12 was originally shown to be required for Integrator function in Drosophila^34^, and subsequently shown to interact directly with the INTS1 subunit *in vitro*^17^. Deletion of this domain preserved strong detection of INTS12 in chromatin but largely eliminated interaction with other Integrator subunits, RNAPII, NELF, and DSIF. In contrast, loss of PHD finger did not affect chromatin association nor interaction with these proteins (Figure 3B, compare lane 7 with lanes 6 and 8; See also Supplementary Figure S3D, lower panel; and heatmap on the right).

The data above show that deletion of the PHD finger has no discernible effects on the interaction of INTS12 with Integrator subunits or RNAPII, suggesting that the PHD finger is not integral to the interaction of INTS12/Integrator with the paused RNAPII complex. Interestingly, however, close comparison of the interactomes of WT INTS12 and INTS12-ΔPHD showed that, while interactions with Integrator, RNAPII and DSIF were unaffected by the loss of the PHD, all subunits of the NELF complex in fact showed slightly reduced Integrator interaction in the ΔPHD mutant (Figure 3D).

Several reports previously demonstrated a direct association of Integrator and NELF^35,36,17^ and through inspection of published *in vitro* cross-linking mass spectrometry data^17^, we noticed an extensive network of interactions, specifically between INTS12 and the NELF-A subunit of NELF. Based on this tentative information, we immunoprecipitated NELF-A from chromatin extracts, and carried out proteomic analysis of its interactors. Notably, Integrator subunits were the most enriched interactors of NELF-A, even more so than RNAPII subunits (Figure 3E). INTS12 was the most enriched among the Integrator subunits co-precipitated with NELF-A, further supporting the idea of an interaction between INTS12 and NELF-A (Figure 3E and 3F).

### The INTS12 PHD finger interacts directly with NELF-A’s C-terminal domain

With the indications above of an interaction between INTS12 and NELF (*cf.* Figure 3), and NELF-A in particular, we now used AlphaFold multimer to model a 3D structure of full-length INTS12 and NELF-A together (Supplementary Figure S4A and B). Strikingly, this identified a high confidence interface between the INTS12 PHD finger and a small (∼10 kDa) ordered domain at the distal C-terminus of the flexible NELF-A tentacle (Figure 4A and B; Supplementary Figure S4A-F), which we hereafter refer to as the NELF-A C-terminal domain (CTD). To corroborate the *in silico* structure prediction, GST-INTS12 PHD and 6xHis tagged NELF-A CTD (amino acids 431-528) were co-expressed in *E. coli* (Supplementary Figure S4G and H). Pull downs of either 6xHis-NELF-A CTD on Nickel-NTA beads or GST-INTS12 PHD on glutathione beads co-precipitated the reciprocal domain, which was not observed in the control lanes, indicating a specific and direct interaction between the two domains *in vitro* (Figure 4C, compare lane 6 with 4-5; and lane 9 with 7-8).

**Figure 4.**
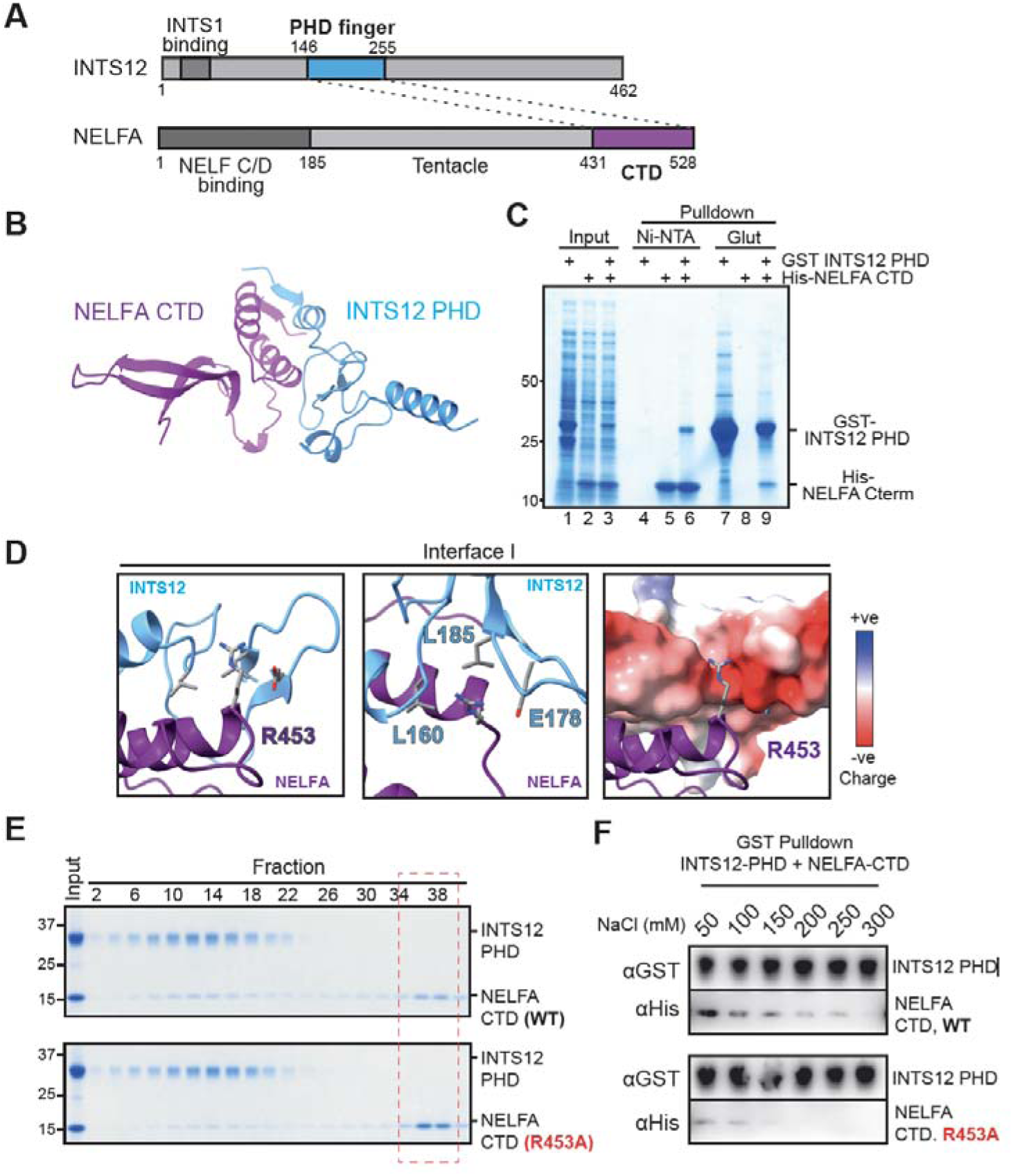
INTS12 PHD interacts with the C-terminal domain of NELFA. **A.** Schematic of the INTS12 and NELFA proteins, showing the interaction identified by structural prediction. Domains with known structure (determined by cryo-EM or ColabFold) are in highlighted in bold, while disordered regions are in light grey. **B.** The INTS12 PHD (145-225) / NELFA CTD (431-528) heterodimer, as predicted by ColabFold. **C.** Instant blue gel staining of pull-downs from lysates from bacteria expressing either individual GST-INTS12 PHD or His-NELFA CTD domains, or lysate with co-expression of the two domains. Pull downs were carried out with either Nickel-NTA (Ni-NTA) or glutathione (Glut) beads. **D.** Close up shots of the INTS12 PHD-NELFA CTD interface, depicting sub-interface I. Residues at the interfaces are shown, labelled in bold and colored according to element. In the right image, the NELFA CTD domain is depicted as ribbon while INTS12 PHD domain is depicted as surface charge. **E.** Instant blue staining of fractions from size-exclusion chromatography of purified GST-INTS12 PHD pre-incubated with either purified NELFA CTD WT or NELFA CTD R453A mutant. Even number fractions with intervals of 4 are labelled. **F.** Western blot of differential salt-washes of GST-INTS12 PHD beads, after incubation and binding with either wild type NELFA CTD (upper) or R453A NELFA CTD (lower).

Through closer examination of the INTS12 PHD–NELF-A CTD interface of the structure predicted by AlphaFold, we identified a striking “lock-and-key-like” insertion of an arginine residue (R_453_) from NELF-A into the substrate recognition site of the INTS12 PHD finger, a pocket defined by several residues, including INTS12 Leucine_160_, Leucine _185_, and Glutamate_178_ (Figure 4D). Indeed, the predicted PHD binding pocket displays high negative charge, consistent with it recognizing a positively charged arginine residue (Figure 4D, right).

This interface resembles that of canonical PHD fingers binding to the H3 tail and H3K4 residue, whether methylated or unmethylated^19–24^, pointing to conservation in the way PHD fingers bind, regardless of the type of binding partner. A series of hydrophobic interactions was also seen at the interface, including with the N-terminal α-helical extension of INTS12 PHD and a hydrophobic pocket in NELF-A (Interface II & III; Supplementary Figure S4F).

The NELF-A CTD is highly conserved between human, *Xenopus laevis* and *Drosophila melanogaster* (Supplementary Figure S5A). Indeed, in the equivalent Drosophila (D.m.) structure prediction, high structural conservation of the interaction was observed, and several residues at the interface were conserved at the sequence level, including at NELF-A Arginine_453_ (R_453_) (D.m. R_1175_) and INTS12 L_160_ (D.m. L_129_) and E_178_ (D.m. E_147_) (Supplementary Figure S5B and S5C). The remaining hydrophobic interfaces appeared to be less well conserved in Drosophila, leading us to hypothesize that recognition of NELF-A R_453_ by the INTS12 PHD finger might be most critical to the interaction.

With this in mind, we investigated the importance of the NELF-A R_453_ residue for the INTS12 PHD – NELF-A CTD interaction using *in vitro* approaches. The pure GST-INTS12 PHD and 6xHis-NELF-A CTD (WT or R453A) (Supplementary Figure S4G and S4H) were incubated together at equimolar concentrations in the presence of 150 mM NaCl, before fractionation by size exclusion chromatography (Figure 4E). A significant fraction of the mixed proteins co-eluted (Figure 4E, upper, fractions 8-18), validating a direct interaction. Importantly, however, a fraction of unbound NELF-A CTD was also visible eluting in the later fractions (Figure 4E, upper, fractions 34-40 (red stippled box)), suggesting that the interaction is of relatively low affinity. Importantly, incubation of INTS12 PHD with the NELF-A CTD R453A mutant showed a weakened interaction, evident by the relative accumulation of unbound, mutant NELF-A CTD in the later eluting fractions (Figure 4E, lower (red stippled box); and Supplementary Figure S6A). Although the interaction was weakened, it was not completely abolished, as some mutant NELF-A CTD still eluted in fractions 6-18 with INTS12 PHD. In an analogous assay conducted at a lower salt concentration (50 mM NaCl), enhanced binding was observed, shown by an almost complete absence of unbound NELF-A CTD, and with a clear reduction in binding in the R453A mutant (Supplementary Figure S6B and S6C). We also assessed how R453A affects the interaction using an *in vitro* pull-down assay. Here, the purified WT or mutant NELF CTD were incubated with INTS12 PHD on glutathione beads, resulting in association of the NELF-A CTD with the GST-tagged INTS12 PHD domain (Figure 4F). We then washed the beads with increasing concentrations of NaCl and assayed the protein remaining on the beads. INTS12 PHD interaction with WT NELF-A CTD was stable in salt concentrations up to 250 mM, while the R453A mutant was washed away beyond detection at much lower salt concentration. Together, these results again indicate that R453A mutation of the NELF-A CTD destabilizes the interaction with INTS12 PHD, validating the AlphaFold structural prediction and pointing to the potential importance of this residue *in vivo*.

### Evidence for competition between INTS12 / NELF-A microdomains *in vivo*

We now studied the function of NELF-A and its CTD in cells. We first determined whether the NELF-A CTD is required for the integrity of the NELF complex and its interaction with RNAPII, Integrator, and other proteins by IP-mass spec analysis. Comparison of the NELF-A WT and ΔCTD interactomes indicated that the mutant NELF complexes retain structural integrity and bind both RNAPII and Integrator (Figure 5A), indicating that the NELF-A CTD plays little or no role in NELF complex assembly or overall partner binding at the promoter-proximal pause, similarly to INTS12’s PHD (cf. Figure 3B and 3C). Importantly, INTS12 remained an enriched Integrator subunit upon deletion of the NELF-A CTD, indicating that, as expected, interactions between NELF and Integrator beyond those mediated by NELF-A’s CTD and the INTS12 PHD are sufficient to support overall NELF-Integrator interaction.

**Figure 5.**
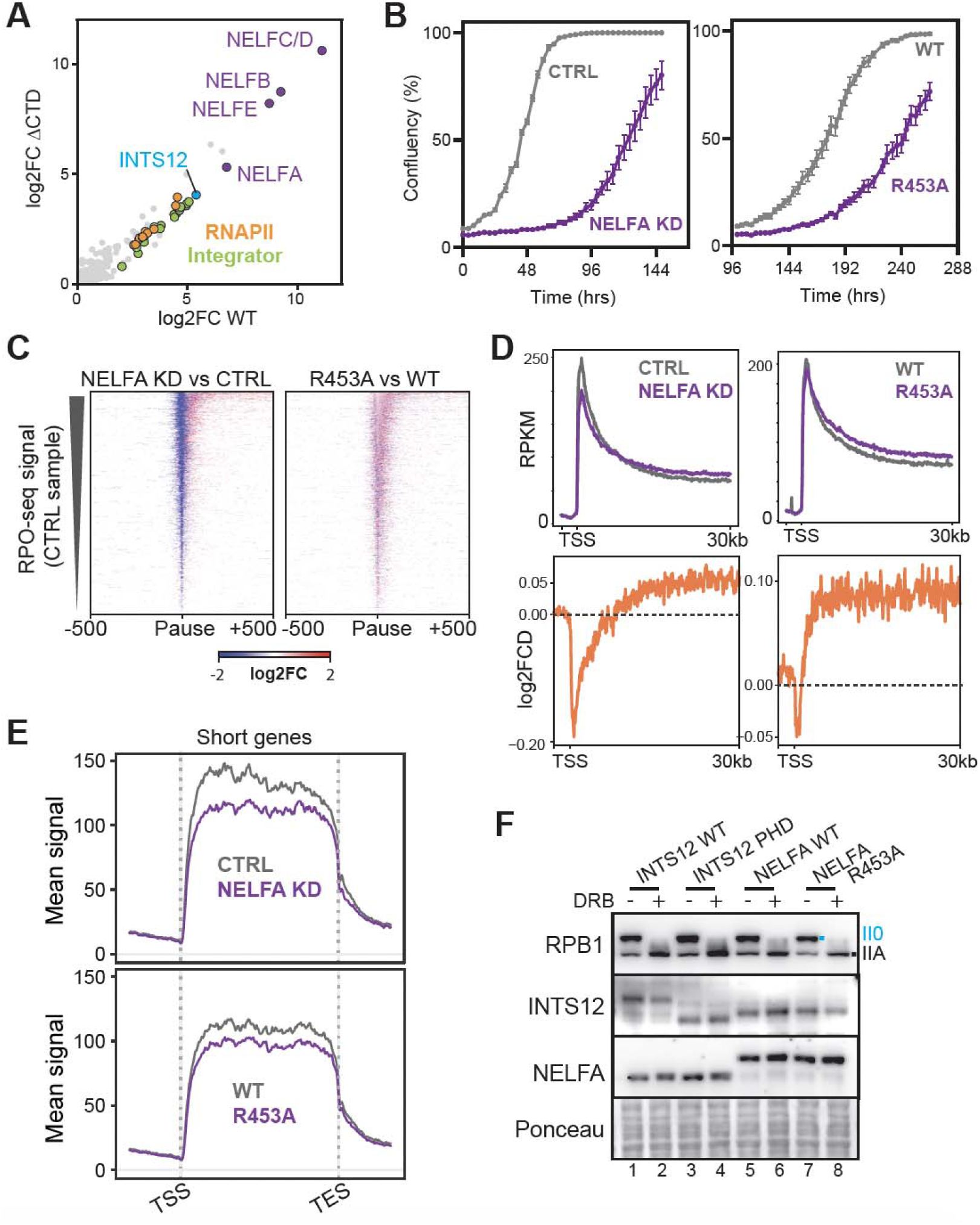
Transcriptional effects of NELFA depletion and NELFA R453A mutation. **A.** Proteomic analysis of FLAG-NELFA chromatin IP fractions. Interactomes of NELFA WT and NELFA CTD are plotted together for comparison. Data is shown as log2 fold-change of NELF pull downs vs negative control (non-FLAG parental cells). Subunits of NELF, RNAPII and Integrator complexes are highlighted. **B.** Growth of cell lines where endogenous NELFA was knocked down (left) or replaced with exogenously expressed WT or R453A mutant NELFA (right). **C.** Average log2 fold-change in PRO-seq signal between NELFA KD/CTRL and NELFA WT/NELFA R453A. Note the loss of normal pausing site position (blue) in cells losing NELFA, which is not occurring in R453A cells. **D. Upper,** Average TT_chem_-seq signal across the first 30kb of protein coding genes, aligned at transcription-start sites (TSS). Comparisons for NELFA knockdown (CTRL vs NELFA KD) and expression of NELFA WT and R453A mutant forms in the switchover systems are shown. **Lower,** Average log2 fold-change of TTchem-seq signals for NELFA/CTRL and WT/R453A. **E.** Average TT_chem_-seq signal across a subset of short protein coding genes (<14kb). **F.** Western blots analysis following switchover of endogenous INTS12 or NELF with INTS12 WT & ΔPHD and NELFA WT & R453A, respectively, with or without DRB treatment. See also Supplementary Figure S7 and S8.

We next generated cell lines containing constructs expressing either WT NELF-A or NELF-A with the R453A mutation. These were NELF-A siRNA resistant constructs, enabling siRNA knock-down of endogenous NELF-A (Supplementary Figure S7A, upper) to test the functional properties of exogenously encoded proteins expressed at similar levels (Supplementary Figure S7A, lower). Importantly, cells expressing NELFA R453A were compromised for growth (Figure 5B), pointing to a crucial role in transcription.

Having determined that disruption of the NELF-A CTD does not appear to affect the overall interactions of NELF, we investigated the effect of NELF mutation on the characteristics of promoter-proximal pausing. Previous results from the Shilatifard laboratory showed that upon rapid loss of NELF, RNAPII pauses downstream from the canonical pause site, in the +1 nucleosomal dyad-associated region^37^. Our PRO-Seq analysis^38^ showed that depletion of NELF-A as expected results in decreased pausing at the canonical pause site and increased movement of RNAPII pausing downstream, but that, interestingly, point mutation of NELF-A (R453A) had no such effect; the site of pausing appeared largely unchanged relative to the wild type control (Figure 5C; see also Supplementary Figure S8). Together with the proteomic analysis above, this supports the idea that the interaction between the INTS12 PHD and the CTD of NELF-A is not required for assembly or positioning of pausing complexes.

We then carried out TT_chem_-seq analyses of transcription to compare the effect of NELFA knockdown with the mutation of R453A that selectively diminishes interaction with INTS12. Depletion of NELF-A by siRNA knockdown resulted in a relative decrease in reads in the 5’ ends of genes, followed by more reads in gene bodies (Figure 5D, Left, Upper; see also Supplementary Figure S7B). These effects on transcription are consistent with previous findings that NELF-A depletion results in loss of RNAPII occupancy at the 5’ ends of protein-coding genes by eNET-seq^26^. Importantly, the NELF-A R453A mutant closely recapitulated NELFA knockdown, pointing to the importance of INTS12 interaction for NELF function (Figure 5D, Right; see also Supplementary Figure S7B-S7D). These changes were particularly clear from difference maps (Figure 5D, Lower). Similarly, short genes were relatively *less* transcribed in both R453A cells and upon NELF-A knockdown (Figure 5E). Hence, although the interactome data and PRO-Seq analysis indicate that NELF complexes containing NELFA R453A form pausing complexes with RNAPII, Integrator and DSIF at the promoter-proximal pause (Figure 5A and C), this single point-mutation nevertheless results in a clear loss of NELF function, manifested by reduced proximal transcription and elevated distal transcription, and penetrant enough to cause significantly compromised cell growth. Notably, the effects of NELFA KD and R453A mutation on nascent RNA synthesis are the opposite of the effects of INTS12 KD or INTS12 PHD mutation (*cf.* Figure 2).

We also used Western blotting of RNAPII to shed light on events at the promoter-proximal pause. As observed previously for INTS12 depletion (*cf.* Figure 1G), introduction of INTS12 ΔPHD resulted in elevated RNAPIIA levels compared with WT (Figure 5F, compare lanes 2 and 4). In apparent accordance with the results above, NELF R453A had the contrary effect of reducing RNAPII levels in the presence of DRB (Figure 5F, compare lanes 6 and 8), consistent with reduced dwell times at the promoter-proximal pause upon loss of NELF function.

Together, these results indicate that although the NELF complex is present at the promoter-proximal pause keeping RNAPII in the canonical pause position, its function in suppressing pause release is greatly disturbed by a single NELF-A R453A point-mutation: the characteristics of RNAPII transcription resembles those of cells in which NELF is absent, and the mutation greatly affects cell growth.

## Discussion

In this study, we show that, in addition to the well-established interactions between NELF-<u>B</u> and INTS<u>6</u> previously identified in the seminal structures of Integrator bound to the NELF-DSIF-RNAPII pausing complex^17,18^, the PHD domain of the intrinsically disordered INTS12 subunit and the flexible C-terminal domain (CTD) of NELF-A mediate a second functional critical NELF-Integrator interaction. This new INTS12/NELF-A interaction is essential for normal transcriptional pausing and pause release. Disrupting this interaction has distinct consequences depending on whether NELF-A or INTS12 is mutated, suggesting that this interface can compete with other molecular interactions and thereby modulate transcriptional pausing and pause release.

### A PHD finger in Integrator complex that recognizes a non-histone protein ligand

While the overall architecture of the 1.6 MDa Integrator complex and most of its 14–15 individual subunits has been characterized^17,18^, the structure and function of the largely disordered INTS12 subunit have remained poorly understood. Our data demonstrate that, despite having modest effect on Integrator integrety and on assembly of the RNAPII-NELF-DSIF-Integrator complex at the promoter-proximal pause, loss of human INTS12 and its PHD results in transcriptional defects resembling the well-established consequences of Integrator loss-of-function at protein-coding genes. These defects include increased pausing at the 5’ end, decreased premature termination, impaired transcript elongation, and elevated expression of certain short genes^37,39,30,27,25^. We also observed that INTS12 deficiency leads to dramatically reduced elongation rates—a phenomenon that, to our knowledge, has not previously been reported in the context of Integrator mutation or loss.

These findings are consistent with previous conclusions that Integrator targets immature RNAPII complexes for termination; thus, disruption of Integrator function results in increased pause dwell times and release of incompetent RNAPII elongation complexes into gene bodies, compromising transcript elongation^30,1,2^. In contrast to its role at protein-coding genes, INTS12 loss has only minor effects on snRNA processing, differing from a study on Drosophila Integrator^34^, which reported that a domain in the N-terminus of INTS12 is required for normal snRNA processing, while the PHD is dispensable. This discrepancy remains unclear, but one possibility is that the earlier study, conducted prior to the discovery of Integrator’s role at protein-coding genes, primarily focused on a relatively mild snRNA processing phenotype in cells lacking the INTS12 N-terminus. This phenotype might thus result from indirect disruption of other Integrator components, such as INTS1 or the cleavage module (INTS4, INTS9, and INTS11), which are known to be essential for snRNA processing^33^.

We identify a direct interaction between the PHD finger of INTS12 and a short CTD segment of NELF-A. Structurally, this interaction closely mimics previously characterized PHDs binding to histone H3K4 residues^23^, with the INTS12 PHD finger specifically “recognizing” R453 in the human NELF-A CTD. Although we initially hypothesized that the INTS12 PHD finger would interact with histone H3, a screen of 123 annotated human PHD fingers revealed that only a minority display histone-binding activity, and INTS12’a PHD does not belong in this subset^24^. Instead, our identification of an alternative binding mode for the PHD finger further supports the notion that PHD fingers are not restricted to histone binding^40^. Consistent with this, recent cryo-EM study indeed also concluded that a direct interaction between Integrator and the nucleosome seems unlikely^18^.

### Microdomains in flexible tethers

While great progress has been made in understanding protein-protein interactions through the revolution in Cryo-EM analysis^41^, the interest in so-called intrinsically disordered regions (IDRs) has concurrently grown dramatically. Intriguingly, up to 50% of the human proteome consists of disordered sequences, which are especially enriched among proteins that regulate chromatin and transcription^42^. Both INTS12 and NELF-A are dominated by IDRs, with the interaction we describe being between short microdomains. The NELF-A CTD segment interacting with INTS12 conforms to the general properties of short linear motif (SLiM), expect that it binds to another more structured microdomain that is also position in a long, disordered tether. This configuration, with two interaction motifs placed as satellites in long tethers, seems ideally suited for association only after being placed in vicinity of each other by the stable interaction of DSIF, NELF, and Integrator in the pausing complex. We speculate that the association of the two domains is diffusion-limited and therefore may contribute to a proof-reading step of pausing complex assembly. In this model, longer dwell increases the likelihood of these two domains interacting, leading to Integrator activation, and eventually premature transcription termination (Figure 6B, right).

**Figure 6.**
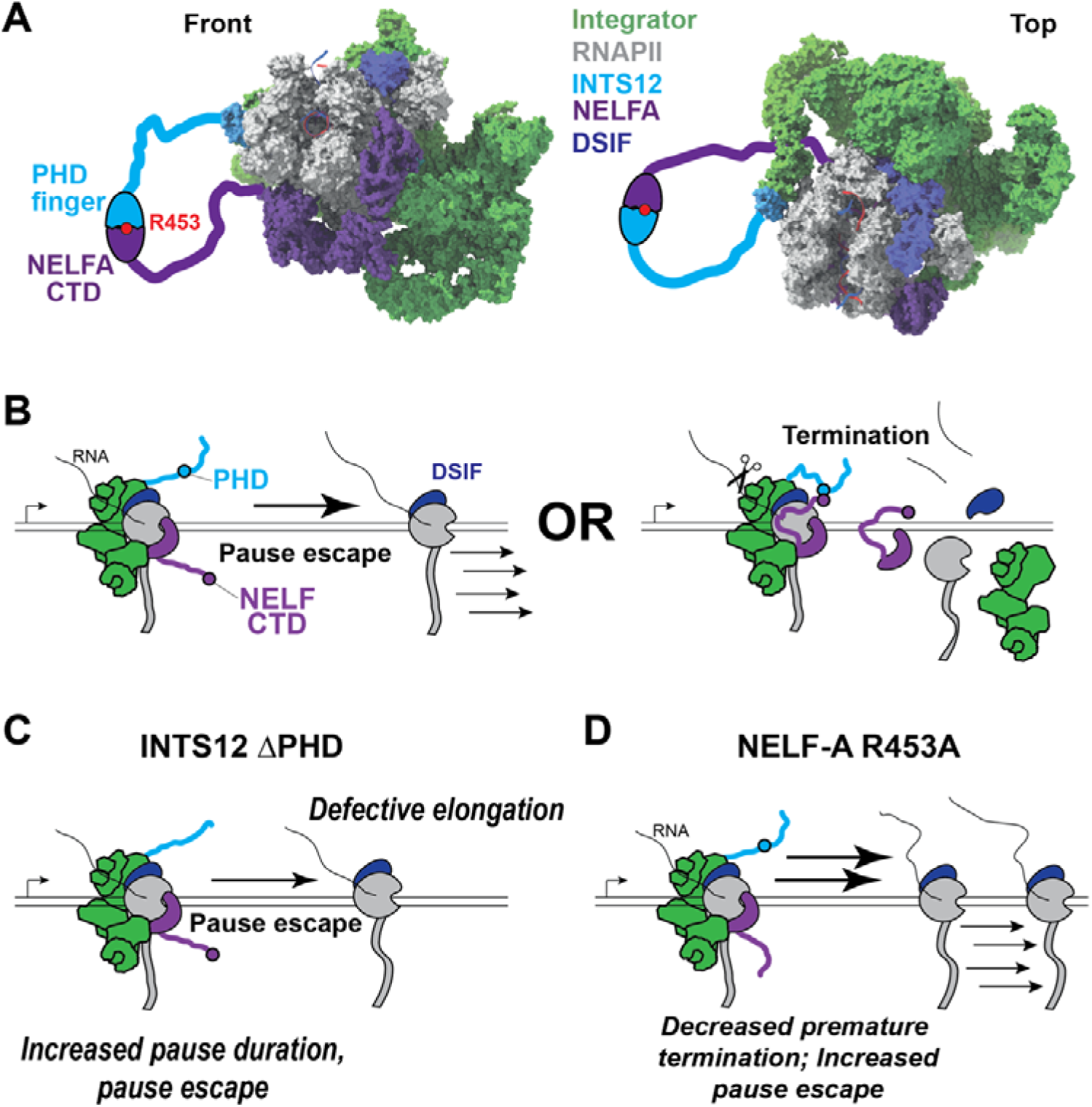
A working model for the role of INTS12 and NELFA during promoter-proximal pausing and release. **A.** Two angles of a model depicting the flexible ‘tentacles’ of INTS12 containing the PHD (light blue bold line) and NELFA’s CTD (purple bold line) superimposed on the cryo-EM structure of the RNAPII-DSIF-NELF-Integrator complex (PDB: 7PKS). In this model, a relatively low affinity of the PHD-CTD interaction requires prior binding of Integrator and NELF with DSIF and RNAPII in the promotor-proximal pausing complex for domain association. NELF-A Arginine 453 is critical for stable PHD-CTD interaction. **B.** The competitive binding of the INTS12 PHD with unknown proteins to help trigger pause release (left), or with NELF-A’s CTD to help trigger premature termination (right). **C.** When the PHD is missing, INTS12 neither interacts with positive transcription factors of unknown origin, nor with NELF-A’s CTD, which results in longer pause dwell times, and in defective elongation complexes being released from pause sites. **D.** When NELF-A’s CTD is mutated, premature termination is decreased and more RNAPII elongation complexes escape the pause.

The downstream consequence of successful INTS12 PHD–NELF-A CTD interaction remains unknown. We do not detect major defects in Integrator phosphatase activity upon INTS12 depletion, but further investigations are needed to explore other possible mechanisms involving conformational activation of Integrator’s RNA cleavage activity, or recruitment of unknown factors that are necessary for RNAPII removal. Others have proposed that premature termination at the promoter-proximal pause may require remodeling of the DSIF upstream DNA clamp, which might in turn be an important pre-requisite for unloading RNAPII^18^. Whether and how the INTS12/NELF interaction contributes to this remains to be investigated.

### A revised model of Integrator at promoter-proximal pause

Current models depict Integrator as a transiently recruited termination factor that acts on a preformed RNAPII, DSIF, and NELF pausing complex, de-phosphorylating the CTD, and utilizing INTS11-mediated nascent RNA cleavage to trigger early termination^1,14,2,15^. Our findings, together with numerous high-resolution structures revealing all of the above proteins in a stable complex, instead support a model in which Integrator is an integral, constitutive component of the pausing^17,18^. In this configuration, RNAPII, DSIF, NELF and Integrator form a single stable assembly at the pause, interconnected through multiple protein-proteins and protein-RNA interactions, including the INTS12 PHD/NELF-A CTD interface described here. This view of Integrator and promoter-proximal pausing alters our interpretation of the impact that Integrator mutations have on transcription. In mutants with loss of Integrator, the remaining RNAPII pausing complex based on DSIF and NELF interactions may thus only support incomplete, suboptimal pausing, while loss of NELF may still allow pausing by an Integrator-DSIF-RNAPII complex, which could help explain the surprisingly modest global effects of NELF loss on transcription^37,26^. In this model, the increase in promoter-escape in mutants with Integrator loss thus reflects not only a decrease in premature RNAPII termination, but also a dysfunctional pausing complex that results in increased pause dwell times and concurrently allows defective RNAPII elongation complexes to escape the promoter. Distinguishing defects in premature RNAPII termination from dysfunctional pausing complexes represents an interesting future challenge.

We propose that under normal conditions, RNAPII is associated with Integrator, NELF and DSIF at the pause, and that interaction between the INTS12 PHD and NELF-A’s CTD specifically regulates whether to release or terminate the RNAPII complex. RNAPII occupancy remains at the canonical pause position in NELF-A R435A cells, indicating that the structural integrity of the pausing complex is maintained. Nevertheless, this mutation perturbs transcription and viability to an extent that mimics complete loss of the NELF-A. Indeed, based on the essential role of the R453 residue and its surprising lack of effect on establishment of the pausing complex, it is tempting to speculate that the indispensable function of NELF may lie not in its ability to establish this complex, but in its interaction with Integrator and the PHD domain of INTS12.

Together, these results support a model in which small, regulated microdomain embedded within disordered tethers fine-tune the behavior of a constitutive RNAPII-DSIF-NELF-Integrator pausing complex. The INTS12 PHD/NELF-A CTD interaction appear as a key determinant of whether paused RNAPII is efficiently terminated or released into productive elongation. Further investigation is need to understand the direct downstream effect of this interaction between the two microdomains and if/which other interfaces can disturb or alter this interaction.

## Acknowledgements

This work was supported by a Laureate grant from the Novo Nordisk Foundation (NNF19OC0055875), a Chair grant (DNRF153) and a Center of Excellence grant (DNRF166) from the Danish National Research Foundation to JQS, and by a fellowship from the Danish Cancer Society to J.L. It was also funded by the Francis Crick Institute, which receives funding from Cancer Research UK [FC001166], the UK Medical Research Council [FC001166], and the Wellcome Trust [FC001166]. Mass spectrometry analyses were performed by the Proteomics Research Infrastructure at the University of Copenhagen (UCPH), supported by the Novo Nordisk Foundation (NNF) (grant agreement number NNF19SA0059305). Work in the Bentley lab was supported by NIH grant R35 GM144336. R.S. is supported by the UC Denver RNA Bioscience Initiative. We thank Eric Wagner, Prashanth Bajpe and others for gifts of plasmids. Advanced sequencing facility and peptide chemistry platforms at the Francis Crick Institute for expert assistance. Svejstrup lab members are thanked for helpful input and discussions.

## Author contributions

DBl and JQS conceived the study. DBl performed most of the experiments with contributions from PB, PD, and RYS. DLM did protein purification and size-exclusion chromatography. NF prepared eNET-seq libraries. RM and RS performed bioinformatic analysis. DBl and JQS wrote the manuscript, with important input from DB and ABDS. DB and JQS guided and obtained funding for the project.

## Materials and Methods

### Data and code availability

Genome-wide data that support the findings of this study have been deposited in GEO (https://www.ncbi.nlm.nih.gov/geo/) with the accession codes GSE318641; GSE268976; GSE253353. Any additional information required to reanalyze the data reported in this paper is available from the lead contact upon request.

### Cell lines and culture conditions

HEK293 Flp-In TREx cells were cultured in DMEM (high glucose, GlutaMAX, pyruvate) supplemented with 10% FBS, 100 U/ml penicillin and 100 ug/ml streptomycin. Cells were cultured with 5% CO_2_ and routinely passaged 2x per week. Stable cell lines were made by integration of respective constructs into the Flp-In locus of HEK293 Flp-In TREx cells. Parental HEK293 Flp-In TREx cells were transfected with a 19:1 mixture of pOG44 and the respective vector containing FRT sites and selected for integration in 100ug/ml hygromycin. Polyclonal cell pools were used as the respective cell line for further experiments. INTS12 KO cells were made by first generating a stable cell line with expression of sgRNA resistant INTS12. For knock-out of endogenous INTS12, cells were transfected with pX459 INTS12 and selected for several days in 2ug/ml puromycin. Single cells were then seeded from the transfected cell pool and clones were picked and screened for INTS12 KO by western blot. Two single clones were used for initial characterization and clone #1 was used for subsequent experiments.

### Plasmid construction

pX495_INTS12 was made by first annealing oligos containing the INTS12 targeting sgRNA construct and then ligating the annealed duplex into a BbsI digested pX459 plasmid backbone. pDEST_FLAG/HA_TO_INTS12_siRES for generation of cells with inducible expression of siRNA resistant INTS12 was made by gateway cloning. INTS12 was amplified with flanking attB sites from a human cDNA library. The attB INTS12 fragment was cloned into the pDONR2223 donor vector using BP clonase and subsequently into the pDEST_FLAG/HA_TO destination vector using LR clonase. The INTS12 cDNA was then made resistant to INTS12 siRNA #2 by site directed mutagenesis. Note that the INTS12 cDNA was already resistant to INTS12 siRNA #3 as this sequence lies in the INTS12 3’ UTR. Hence, endogenous INTS12 could then be knocked down using a combination of siRNA #2 and #3. Subsequent deletions of the PHD finger domain and N-terminus were then made by standard mutagenesis. For construction of pcDNA5_FLAG/HA_TO_NELF-A_siRES, NELF-A was cloned along with flanking EcoRI and BamHI sites and FLAG/HA fragment from pDEST_FLAG/HA_TO and the fragments were Gibson assembled into the pcDNA5_TO backbone after cutting with EcoRI and BamHI. Constructs were then made resistant to NELF-A siRNA #2 and #4, however only siRNA #4 was used in experiments. Subsequent CTD deletion and R453A mutation was then made by standard mutagenesis. pGEX4T1 and pET28a vectors were made by Gibson cloning respective INTS12 and NELF-A fragments into the relevant plasmid backbone.

### Cell growth assays

For cell growth assays, 5000 cells were seeded in a 96-well plate. The growth was monitored using an IncuCyte S3 Live-Cell Analysis System (Sartorius). Images of live cells were captured every 3 or 4 hours for up to 5 days, generally until cells reached 100% confluency. Images were analyzed in the IncuCuyte S3 image analysis software, with an output of cell confluency (%).

For clonogenic survival assays, 1000 cells were seeded into 35cm dishes and left to grow for approximately 10 days to allow formation of colonies. Cells were washed in PBS and fixed in 4% formaldehyde for 15mins at RT. Cells were washed 2x in PBS and stained with crystal violet solution for 30mins at RT. Cells were washed 3x in PBS to remove all excess crystal violet and dried overnight before imaging.

### Cell lysate preparation and cell fractionation

Human whole cell lysates were prepared by resuspending cells in lysis buffer (20mM TRIS-HCl pH 7.5; 150mM NaOH; 1.5mM MgCl_2_; 10% glycerol; 250U/ml Basemuncher; protease & phosphatase inhibitors) and rotating for 1hr @ 4°C. Samples were centrifuged 20mins; max speed to pellet debris and supernatant was taken as whole cell lysate.

For cell fractionation cell pellets were resuspended in SE buffer (20mM HEPES-NaOH pH=7.5; 150mM KAc; 1.5mM MgCl_2_; 0.5% NP40; 10% glycerol; protease and phosphatase inhibitors fresh) and incubated 15mins on ice. Permeabilized nuclei were pelleted 15mins; 20,000G and the supernatant taken as the soluble fraction. The nuclear pellet was resuspended in 2x pellet volumes CD buffer (20mM HEPES-NaOH pH=7.5; 150mM NaCl; 1.5mM MgCl_2_; 0.05% NP40; 10% glycerol; protease and phosphatase inhibitors fresh) plus 250 U/ml Basemuncher and incubated on a rotating wheel for 1hr at 4°C to digest nucleic acids and solubilize chromatin. Nuclei were pelleted 15mins; 20,000G and the supernatant was taken as the Chrom1 fraction. Chrom1 fractions alone were used for immunoprecipitation purposes, where the volume of CD buffer was adjusted based on the desired volume). For Western blots, the NaCl concentration was adjusted to 500mM to further solubilize chromatin bound proteins in Chrom1 fractions. After incubation for 20mins on ice, samples were centrifuged for 15min @ max speed and supernatant collected as the high salt chromatin fraction.

### Immunoprecipitation

All immunoprecipitations were carried out from Chrom1 fractions prepared using the above approach. Extract was incubated with either Anti-FLAG M2 beads (Sigma) for 2-3hrs rotating at 4°C. Beads were then washed 4x in CD buffer. For proteomics analysis, beads were washed 2x times in TBS to remove detergent and transferred to a clean 1.5ml tube. For western blot, beads were boiled in Laemmli buffer.

### Western blot analysis

Proteins were separated on 4%–15% Criterion TGX, (Biorad)transferred to nitrocellulose membrane and blocked for 1hr in blocking buffer (5% milk+PBST) at room temperature. Membranes were incubated withprimary antibodies in blocking buffer overnight at 4°C. Membranes were washed several times in PBST, incubated with HRP-conjugated secondary antibody in blocking buffer for 1hr min at room temperature, and washed several times in PBST. HRP signal detection was obtained using either SuperSignal West Pico PLUS (Thermo Fisher Scientific, 34580) or Radiance plus (Azure Biosystems, AC2103) ECL reagents, visualization by Amersham imager.

### Mass spectrometry

IP samples for proteomic analysis were carried out in triplicate and samples were prepared in parallel from the point of cell seeding. Washed frozen beads were thawed and incubated for 30 min with elution buffer 1 (2 M Urea; 50 mM Tris-HCl pH 7.5; 2 mM DTT; 20 µg/ml trypsin) followed by a second elution for 5 min with elution buffer 2 (2 M Urea; 50 mM Tris-HCl pH 7.5; 10 mM Chloroacetamide). Both eluates were combined and further incubated at room temperature over-night. Tryptic peptide mixtures were acidified to 1% TFA and loaded on Evotips (Evosep). Peptides were separated on 15 cm, 150 μM ID columns packed with C18 beads (1.9 μm) (Pepsep) on an Evosep ONE HPLC applying the ‘30 samples per day’ method, and injected via a CaptiveSpray source and 10 μm emitter into a timsTOF pro mass spectrometer (Bruker) operated in PASEF mode ^43^.

### Pull down from bacterial lysate

pGEX4T-1_INTS12_PHD and pET28a_NELF-A C vectors were either co-expressed or expressed individually in One Shot BL21(DE30) Chemically competent E. coli. Bacteria were grown at 37°C shaking at 220rpm until OD600 of approx. 0.8 was achieved. Expression was induced by addition of 1mM IPTG for 3hrs 30°C shaking at 220rpm. Cells were centrifuged 10mins; 4000G, snap-frozen and stored at - 80°C. Pellets were resuspended in 15ml lysis buffer (20mM TRIS-HCl pH 7.5; 150mM NaCl; 0.05% NP40; 1.5MgCl2; 1mM EDTA; 10uM ZnCl2; 10% Glycerol; Protease inhibitor fresh). Suspensions were sonicated on ice with a tip probe Branson Digital Sonifier 250 with 15% amplitude for a total sonication time of 15mins (15s on / 25s off). Lysates were snap-frozen and stored at −80°C. For pull downs, 1ml lysate was incubated with 25ul bead vol. of either Glutathione Agarose or Ni-NTA beads for 2hrs rotating at 4°C. Beads were washed 4x in lysis buffer and boiled in Laemmli buffer + 50mM DTT for 5mins. The beads were transferred to a Pierce Mini Spin Column and centrifuged 15s; 500G. Samples were separated on 4-15% TGX gels (BioRad) and stained with InstantBlue Coomassie stain (Abcam).

### Protein purification

BL21 cell pellets were resuspended in lysis buffer: 20 mM HEPES-NaOH (pH = 8.0), 150 mM NaCl, 10 % glycerol, 1x Protease Inhibitor Mix (Roche-EDTA), 2 mM - mercaptoethanol complemented with 100 µM ZnCl_2_ for GST-INTS12-PHD or 25 mM imidazole for His-NELFA-CTD and lysed by sonication on ice (5 min, 50 % amplification, 2 s on/2 s off). Lysates were centrifuged at 20000 rpm for 30 min at 4 and pellets were discarded. Clarified lysates were then incubated with appropriate beads (Glutathione sepharose for GST tag (GST-INTS12-PHD) and Ni^2+^-NTA agarose for His tag purification (His-NELFA-CTD)) for 2 hours at 4 rotating. Beads were collected by centrifugation and washed with lysis buffer. Proteins were eluted with lysis buffer complemented with 7 mM Glutathione or 250 mM Imidazole according to the tag: GST or His, respectively. Elutions were further cleaned in the HiTrap Desalting column (Cytiva, Cat #29048684) to remove elution components and reduce the NaCl concentration to 50 mM. Final eluates were aliquoted, and flash frozen in liquid nitrogen before storage at – 80.

### In vitro pulldown with salt wash

GST-INTS12-PHD was mixed with His-NELFA-CTD (WT or R453A) to a final concentration of 4 µM and 6 µM, respectively, in IP buffer: 20 mM HEPES-NaOH (pH = 8.0), 50 mM NaCl, 10 % Glycerol, 0,05 % NP-40, 2 mM-mercaptoethanol, for 1 hr at 30. Then Glutathione Sepharose (Cytiva) was added and incubated for 1 hr at 4. Input was taken and then beads were pelleted at 900 rpm for 2 min. Flow-through was collected from supernatant. Beads were washed with IP buffer with 50 mM NaCl, and the process was repeated with increasing amounts of NaCl: 100, 150, 200, 250 and 300 mM. Beads were collected at each step.

### Size exclusion chromatography

All analytical size exclusion chromatography (SEC) binding assays were conducted on an ÄKTAmicro system (GE Healthcare) using a Superdex 200 Increase 10/300 GL column (Cytiva, cat No. 28-9909-44) equilibrated in 20 mM HEPES-NaOH pH 8.0, 150 mM NaCl and 10 % glycerol. GST-INTS12-PHD (WT) was mixed with His-NELFA-CTD (WT or R453A) in a 1:1 molar ratio (30 µM final concentration) and incubated for 30 minutes at RT in a total volume of 300 µl and spun (5 minutes at 14000 rpm) before injection. The eluted volume was monitored by A260-280, collected in 250 µl fractions and analyzed by SDS-PAGE.

### eNET-seq

eNET-seq experiments were performed as recently described^26^, which - compared with the original NET-seq procedure^44^ - includes optimized MNase (NEB) digestion conditions, RNA decapping with recombinant *S. pombe* Dcp1-Edc1-Dcp2, and incorporation of 12 base UMIs in library construction to permit unambiguous elimination of duplicates^26^.

### TT_chem_-seq

TTchem-seq was performed essentially as described previously^29^. In the INTS12 KO (GSE268976) and INTS12/INTS4 knockdown (GSE253353) experiments, libraries were prepared using the KAPA RNA HyperPrep kit with omission of the fragmentation step. 75 ng of nascent RNA was denatured at 65°C for 5 min. The rest of the procedure was performed following manufacturer’s instructions, with two SPRI bead purifications (Beckman Coulter, A63881) with a bead-to-sample volume ratio of 0.95x and 1x respectively. Libraries were sequenced on Illumina Novaseq with paired-end 100bp reads. For NELF-A knockdown/rescue experiments, 50 ng of enriched RNA was used for library construction with NEBNext Ultra II Directional RNA Library Prep Kit (NEB #E7760L) and NEBNext Multiplex Oligos for Illumina (Unique Dual Index UMI Adaptors RNA Set 1, NEB #E7416L). Libraries were pooled and sequenced single-end 50 on a Nextseq2000 flowcell.

### PRO-seq

Quick PRO-seq (qPRO-seq) libraries were prepared following a previously established protocol^38^ with an optimized cell permeabilization method. Briefly, Flp-In™ T-REx™ 293 cells cultured in a 10 cm dish were trypsinized and washed once with 10 mL of ice-cold PBS. Cells were then permeabilized in 10 mL of Buffer P (10 mM Tris-Cl pH 8.0, 10 mM KCl, 250 mM sucrose, 5 mM MgCl_2_, 1 mM EGTA, 0.1% NP-40, 0.5 mM DTT, 0.05% Tween-20, 10% glycerol, and 0.2 µL/mL RNase inhibitor) for 6 min. After permeabilization, cells were washed once with 10 mL of Buffer W (Buffer P without detergents), pelleted by centrifugation, and resuspended in Buffer Z (10 mM Tris-Cl pH 8.0, 40% glycerol, 5 mM MgCl_2_, 1.1 mM EGTA, 0.5 mM DTT, and 1 µL/mL RNase inhibitor) to a final concentration of 25 million cells/mL. The permeabilization efficiency was verified by trypan blue staining to ensure a rate of at least 90%. Permeabilized cells were flash-frozen in liquid nitrogen and stored at −80°C. For each nuclear run-on reaction, 50 µL (1.25 million) of permeabilized cells were incubated with all four biotin-11-NTPs in a thermomixer at 37°C, 750 RPM for 5 min. Total RNA was extracted using TRIzol LS reagent, and library preparation was performed strictly according to the previous protocol. The final reverse-transcribed products were directly amplified using the following PCR program:95°C for 2 min;5 cycles of (95°C for 30 s, 56°C for 30 s, 72°C for 30 s);14 cycles of (95°C for 30 s, 65°C for 30 s, 72°C for 30 s); final extension at 72°C for 5 min. Amplified products were purified using 1.8x SPRI beads. Libraries were sequenced on an Illumina NextSeq 2000 platform with 66 bp paired-end reads.

### Quantification, bioinformatics and statistical analysis

Throughout this study, analyses were performed using multiple versions of the same bioinformatics tools due to updating of the tools. While minor version differences existed in the software implementations, rigorous validation confirmed that these variations did not significantly impact the key findings or overall conclusions of our research.

### TT_chem_-seq data analysis

Reads were processed using the publicly available nf-core rnseq pipeline v3.3 with the STAR /RSEM^45,46^ option against human genome assembly GRCh38 and Ensembl release 104 transcript annotations. Other options were left as default expect for the TT-seq data which were simultaneously aligned against Saccharomyces cerevisiae R64-1-1 cdna sequences (Ensembl 104) using the “--additional_fasta” flag. For the purposes of visualization, genome alignment BAM files were merged across biological replicates, sorted and indexed using Picard v2.1.1 functions MergeSamFiles, SortSam and BuildBamIndex (all default settings) (http://broadinstitute.github.io/picard). In the case of dataset GSE318299, the 61bp single-end and corresponding separate 11bp UMI raw fastq files, where merged using UMI-tools (v1.1.4) ‘extract’ thereby appending each UMI to the read header (command: umi_tools extract --extract-method=string --bc-pattern=“NNNNNNNNNNN” --stdin=<UMI fastq file>--read2-in=<READ fastq file> --read2-out=<OUTPUT fastq file>). Before read processing, FastqQC (v0.12.1a) and FASTQ Screen (v0.15.3) were used to evaluate the library and read quality of the unprocessed reads. Reads were quality trimmed with Fastp (v0.21.0, default settings except: ‘--trim_poly_x --length_required 25’), then aligned to the hg38.p14 genome assembly guided by gene annotion (ENSEMBL,v112) using STAR (v2.7.9a) in two-pass mode (default settings except: --sjdbGTFfile <GTF_FILE> --outSAMmultNmax 1 --alignEndsType EndToEnd --sjdbGTFtagExonParentTranscript Parent --outSAMstrandField intronMotif --twopassMode Basic --outSAMtype BAM SortedByCoordinate -- outSAMattributes All --outSAMunmapped Within --outFilterIntronMotifs RemoveNoncanonicalUnannotated). Unmapped or multimapping reads were filtered out using samtools ‘view’ (v1.20, settings: -q 255). PCR-duplicates were then eliminated with UMI-tools ‘dedup’. Finally, reads mapping to spliced transcripts (marked with ‘XS’ tags by STAR) were removed using samtools view (settings: ‘-e ![XS]’). Replicate BAM files were merged using SAMtools merge.

Reads Per Kilobase per Million mapped reads (RPKM) scaled BigWig files were generated using Deeptools’ bamCoverage function for each strand using the options “--normalizeUsing RPKM --exactScaling –effectiveGenomeSize (Ramirez et al., 2014). Log fold-change ratio bigwig files were created using deepTools’ bigwigCompare function. Briefly, the genome was partitioned to 50bp bins and per bin coverage calculated separately for the CTRL and treatment samples. A pseudo-count of 1 was added and a new summary signal was calculated as the log2 ratio of each bin in the treatment and CTRL samples. Read depth profiles were created directly from the scaled bigwig files using deepTools v.2.5.3 (Ramirez et al., 2014) using gene definitions from Ensembl release 104 GTF. Metagene profiles were generated using deeptools computeMatrix function with the –scaleregions option for TSS-TES plots or the –reference point with respective gene length cut-off as indicated in respective figures.

### eNET-seq read processing

eNET-seq libraries were sequenced on an Illumina NovaSeq 6000 (2×150). Adapters were trimmed using cutadapt (v2.3) and reads were aligned to the hg38 human genome using Bowtie2 (v2.3.2). PCR duplicates were removed using UMI-tools (v0.5.4) and read coordinates were collapsed to a single base pair coordinate corresponding to the RNA 3’ end. Reads were filtered to only include those with a mapping quality score >=10, and to remove reads that did not align within 5 kb of a protein coding gene, or that aligned to a snoRNA gene. eNET-seq 5’ and 3’ metaplots include all genes longer than 2 kb and separated by >5 kb. In addition, genes had to contain at least one read and the bottom 10% of genes with the lowest signal were excluded. Read counts were normalized by library size and the size of the bin. The mean signal was calculated separately for each biological replicate. The shaded region shows the minimum and maximum mean signal for the replicates, and the center line shows the mean of the replicates.

### eNET-seq read subsampling

eNET-seq read subsampling was performed as previously described^26^. For comparing eNET-seq signal (Fig 1E-F), eNET-seq datasets were down sampled so all libraries being compared had the same number of aligned and filtered reads. To compare RNAPII pausing for different gene regions (Fig 1G), reads were subsampled so each TSS (TSS - +500 bp) or gene body (+500 bp – pAS) region had the same number of aligned/filtered reads for all datasets used in the comparison. Subsampling was performed separately for each individual gene and involved identifying the dataset with the fewest number of reads for the gene region. Reads aligning to the region were then randomly subsampled for the remaining datasets so all samples used in the comparison had the same number of aligned reads for the region.

### eNET-seq; quantification of RNAPII pausing

Pause sites were located as previously described^26^. Pauses were located by identifying positions where eNET-seq signal was >3 standard deviations above the mean for the surrounding 100 bp on each side, there were at least 5 reads at the pause site, and at least 5 additional reads within the window. Pause sites identified within 2 bp of 5’ and 3’ splice sites were excluded from downstream analysis. The fraction of pause reads was calculated by dividing the total number of reads aligned to pause sites by the total number of aligned/filtered reads for the region. Genes were filtered to only include those with at least 1 pause in both the TSS and gene body regions, and a total read density of at least 1 read/kb within the gene body region.

Pausing zones were calculated by identifying the window downstream of the TSS (using 200 bp windows) where the fraction of pause reads was >0.15 and the mean for the pausing zone windows was >0.3. Pausing zones were identified after subsampling reads so each gene had the same number of aligned/filtered reads for all datasets used in the comparison (see subsampling description above). Pausing zones were identified for genes >4 kb in length and separated by >2kb.

### PRO-seq

#### qPRO-seq analysis

Illumina adapters were first trimmed from the 3’ ends of paired-end reads using Cutadapt, with the following parameters: ‘-a GNNNNNNTGGAATTCTCGGGTGCCAAGGAACTCCAGTCAC-A TNNNNNNGATCGTCGGACTGTAGAACTCTGAACGTGT -O 10 --minimum-length 10 - -pair-filter=any’. Unique Molecular Identifiers (UMIs) located at the first 7 bases of both reads were extracted using UMI-tools (’umi_tools extract --quality-filter-mask=10 --quality-encoding=“phred33” --extract-method=string --bc-pattern=NNNNNNN --bc-pattern2=NNNNNNN’). The processed reads were aligned to the hg38 reference genome using Bowtie2 in ‘--very-sensitive’ mode with ‘--non-deterministic --no-discordant --no-unal’ flags. Reads with low mapping quality (MAPQ < 10) were removed via SAMtools. PCR duplicates were then identified and collapsed based on UMIs using the command: ‘umi-tools dedup --paired’. To determine the precise location of active RNAPII, single-base resolution BigWig files were generated from the first base of Read 1 using the bamCoverage function (deepTools) with parameters ‘--samFlagInclude 64 --offset 1 1 --binSize 1’. Forward and reverse strand BigWig files were generated separately using the ‘--filterRNAstrand’ option.

For downstream analysis, human protein-coding genes longer than 2 kb were retrieved from Ensembl (v112). Transcript expression levels were quantified from RNA-seq data of WT Flp-In 293 cells (GSE143537) using Salmon, retaining only the highest-expressed transcript for each gene. To avoid signal interference, genes overlapping with others within a 2 kb window were excluded. Highly paused genes were then identified based on previously reported criteria^11^: Transcripts required ≧2 RPM total PRO-seq coverage within the TSS region (TSS - 100 bp to TSS + 300 bp) in control cells. The position with maximum coverage within this region was defined as the 1st pause peak. Transcripts with ≦ 0.2 RPM at this peak were excluded. In NELF-knockdown cells, the position with maximum coverage within +10 to +500 bp downstream of the 1st pause peak was designated as the 2nd pause peak. To visualize transcription changes around these peaks, heatmaps and metagene profiles were generated using custom python scripts leveraging pyBigWig and matplotlib.

### Structural prediction

Structural predictions were generated using ColabFold^47^ in UCSF ChimeraX version 1.5. Images were rendered in ChimeraX.

## SUPPLEMENTARY FIGURES

**Supplementary Figure S1, related to Figure 1 and 2.**
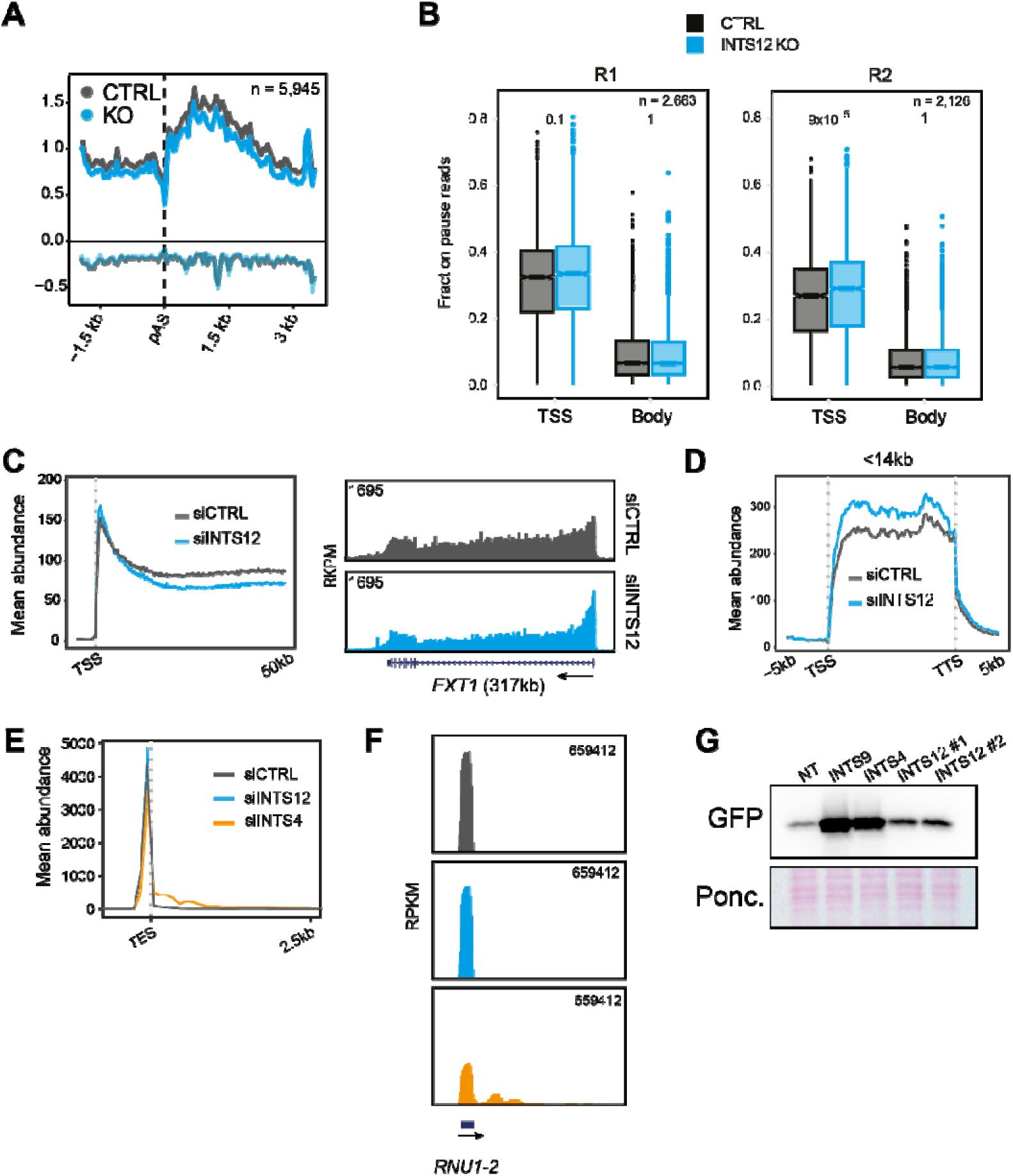
**A.** Average eNET-seq signal at 3’ ends of protein coding genes, aligned on polyA sites. **B.** Box plots showing fraction of paused reads, aka. Pausing propensity, calculated from eNET-seq data (see Materials and Methods for further details). Data for 2x eNET-seq replicates is shown. **C.** Average TTchem-seq signal on protein coding genes in control and INTS12 knockdown conditions. Data is aligned on TSSs. Right: Single gene example for *EXT1.* **D.** Average TTchem-seq signal on a subset of short protein-coding genes in control and INTS12 knockdown conditions. **E.** Average TTchem-seq signal on snRNA genes in INTS12 and INTS4 knockdown, or control cells. **F.** Example of TTchem-seq signal on the *RNU1-2* snRNA gene. **G.** Western blot analysis of whole cell extracts after knockdown of respective target genes and transfection with the human U7 snRNA – GFP reporter construct.

**Supplementary Figure S2, related to Figure 3.**
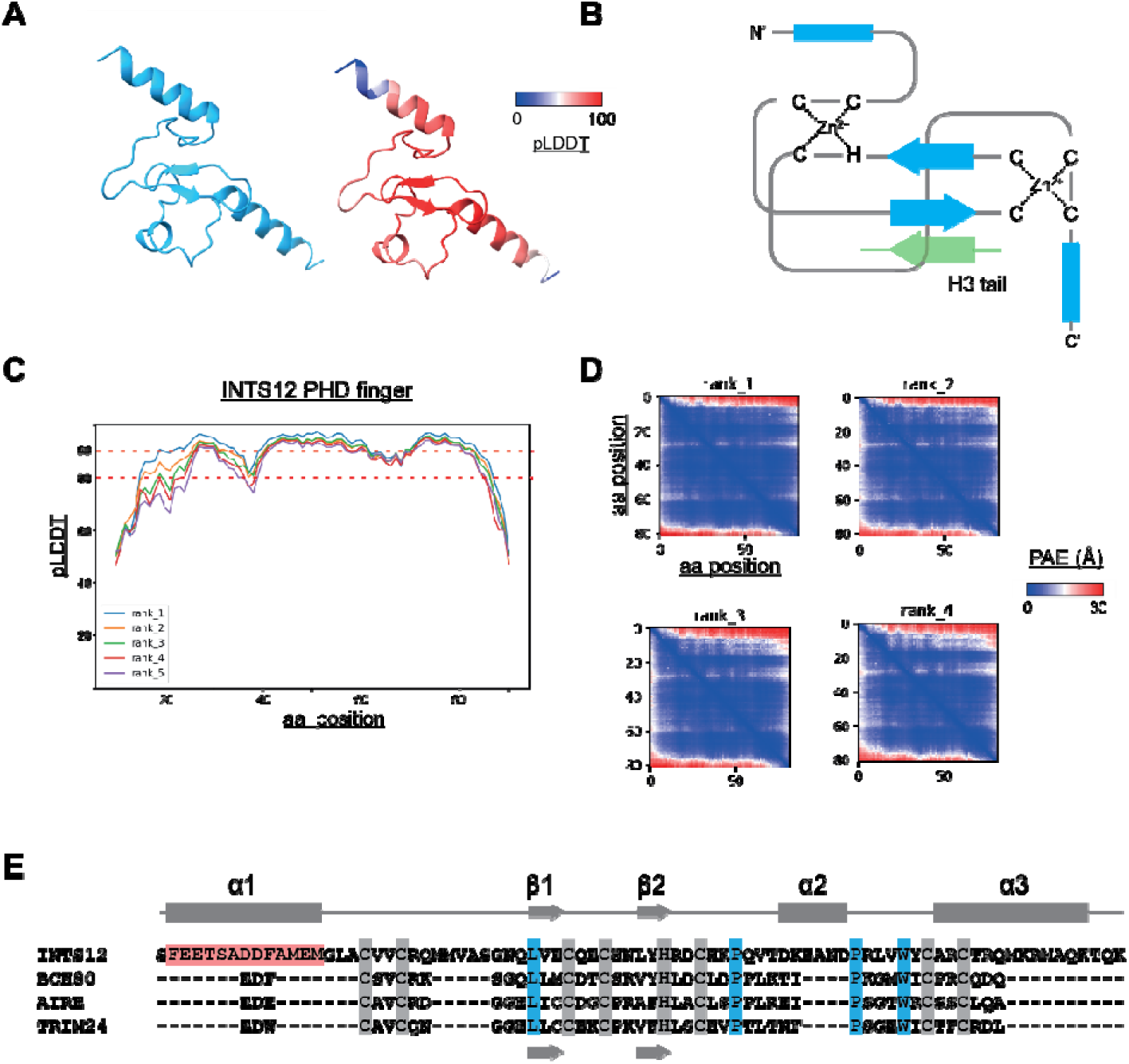
**A.** Structural prediction of the INTS12 PHD finger domain colored in either a single color (left), or according to the predicted local difference distance (pLDDT) score. Red indicates high pLDDT (high confidence) and blue indicates low pLDDT (low confidence). **B.** Schematic representation of the INTS12 PHD finger architecture. The position at which histone H3 N-terminus would putatively bind is also indicated (H3 tail). **C.** pLDDT plots from 5 AlphaFold models for the INTS12 PHD finger domain. Red dotted lines indicate accuracy threshold. **D.** Predicted aligned error (PAE) plots for 4 AlphaFold models, showing confidence in position of residue x relative to residue y. Blue indicates low positional error, measured in Ångstroms (Å). Axes numbers indicate amino acid position in the query sequence. **E.** Alignment of the human INTS12, BCH80, AIRE and TRIM2 PHD fingers. A schematic representation showing predicted secondary structures in the INTS12 PHD finger is shown on top in grey. Meanwhile, grey arrows on the bottom show where the characteristic anti-parallel β-sheets form in structures determined from other PHD fingers. Conserved residues are highlighted, with those residues that co-ordinate Zn^2+^ ions in highlighted in grey. Other conserved residues are highlighted in blue. The unique N-terminal extension is also highlighted in red.

**Supplementary Figure S3, related to Figure 3.**
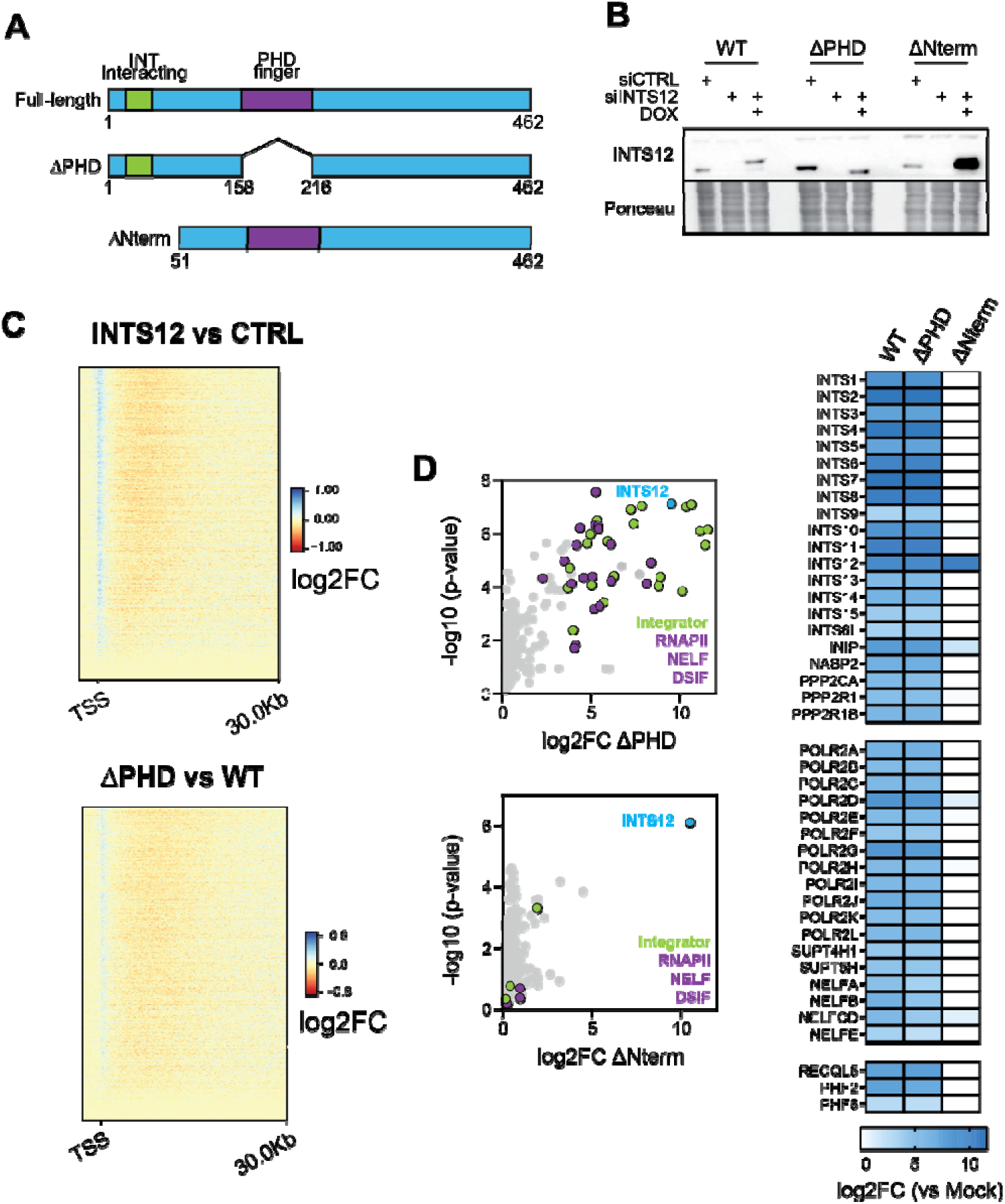
**A.** Schematics of INTS12 constructs expressed in the different switchover cell lines. **B.** Western blot analysis of whole cell lysates from switchover systems, following endogenous INTS12 knockdown and induction of INTS12 WT or ΔPHD transgene expression. **C.** Heatmaps showing log2 fold-change in TTchem-seq signal across the first 30kb of protein-coding genes. Log2FC is given for INTS12 knockdown vs control, and INTS12 ΔPHD vs WT switchovers. Data is aligned at the TSS. **D.** Proteomic analysis of FLAG-INTS12 ΔPHD and -ΔNterm chromatin IP fractions. See Figure 3C for WT control. Left: Scatterplots for the indicated INTS12 forms, showing log2 fold-change relative to a negative control (parental non-FLAG tagged cells). Integrator, RNAPII and DSIF/NELF subunits are highlighted. Right: Heatmap representation of the log2FC values for Integrator, RNAPII pausing complex and several others novel INTS12 interactors.

**Supplementary Figure S4, related to Figure 4.**
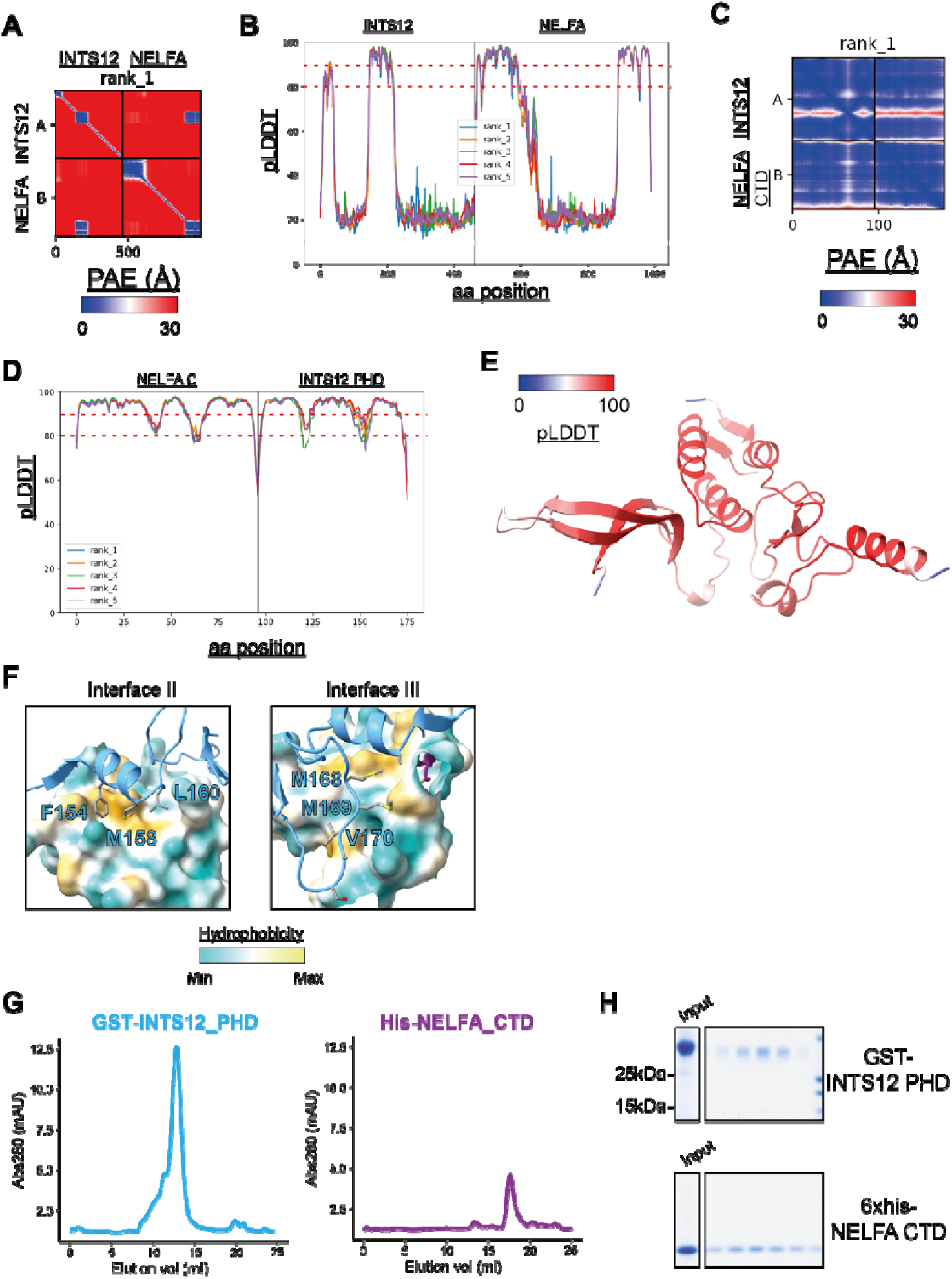
**A.** AlphaFold multimer 2D PAE plots for full-length INTS12 and NELFA amino acid sequences. Number on x axis corresponds to cumulative amino acid position. **B.** pLDDT plots for full-length INTS12 and NELFA proteins. Red dotted lines show confidence thresholds. X axis indicates cumulative amino acid position. **C.** As in A, but for INTS12 PHD and NELFA CTD domains. **D.** As in B, but for INTS12 PHD and NELFA CTD domain prediction. **E.** Predicted structure of INTS12 PHD and NELFA CTD as in Figure 5B but colored by pLDDT score. **F.** Sub-interfaces (II and III). INTS12 PHD is depicted as ribbon, while for NELFA CTD the surface hydrophobicity is shown. **G.** UV absorption of Superdex 200 elutions, with samples containing individual purified INTS12 GST-PHD domain (left) or 6xHis-NELFA CTD domain (right) as inputs. **H.** Instant Blue staining of fractions from G.

**Supplementary Figure S5, related to Figure 4.**
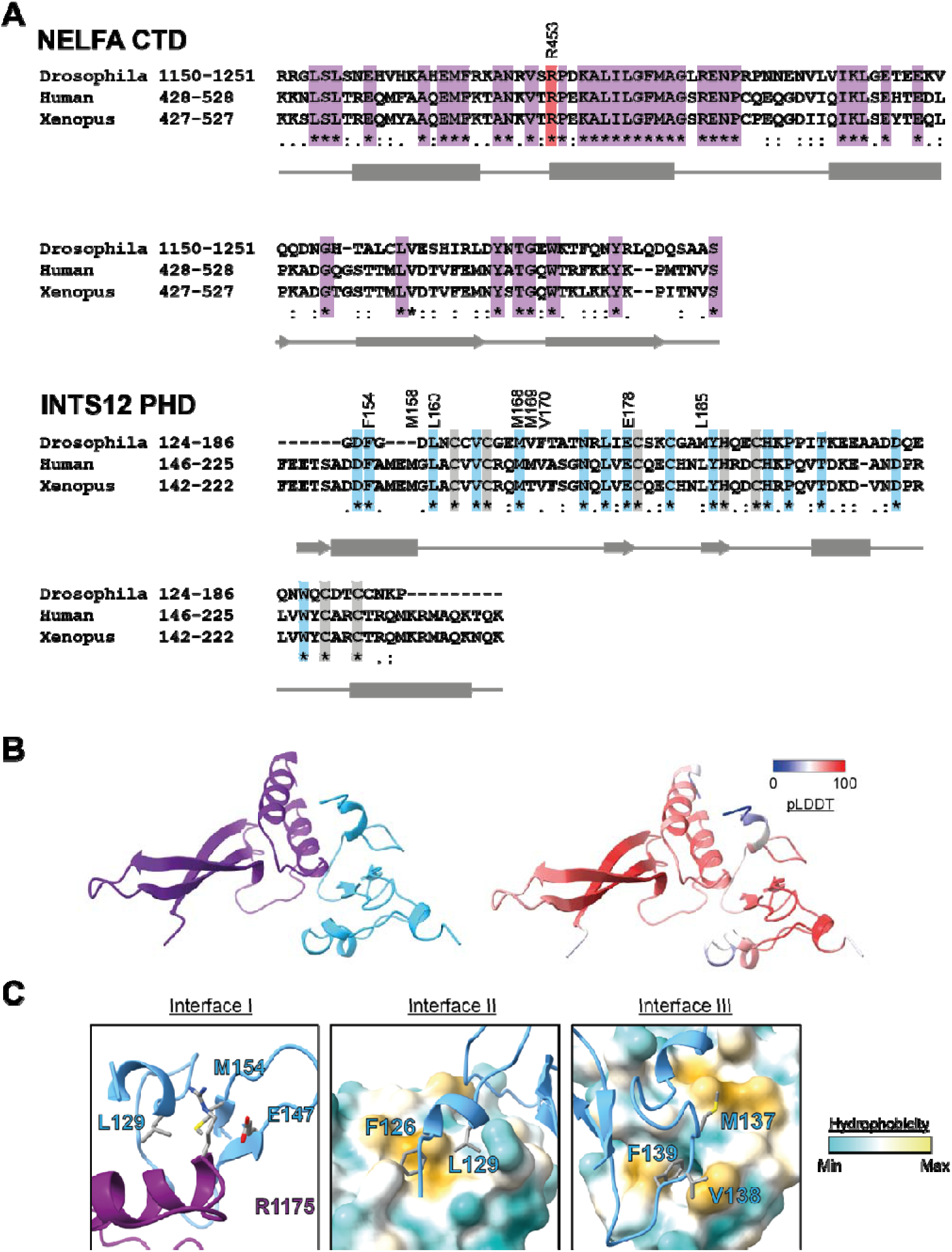
**A.** Alignments of INTS12 PHD finger and NELFA CTD domains from Human, Drosophila and Xenopus. Conserved residues are highlighted in purple or blue for NELFA and INTS12 domains, respectively. Schematics depicting secondary structures are shown below alignments in grey. For INTS12, highly conserved Zn^2+^ coordinating residues are also highlighted in grey. Key residues at the INTS12 PHD / NELFA CTD interface are labelled with the human residue numbers. R453 is highlighted in red. **B.** AlphaFold multimer predictions of the Drosophila INTS12 PHD and NELFA CTD domains. Left: colored by domain. Right: colored by pLDDT score. **C.** Zoomed in shots of the structural interface in the structure predicted from Drosophila INTS12 PHD and NELFA CTD domains. Key residues at the interface are labelled. For Interfaces II and III, the NELFA CTD surface is colored according to hydrophobicity.

**Supplementary Figure S6, related to Figure 4.**
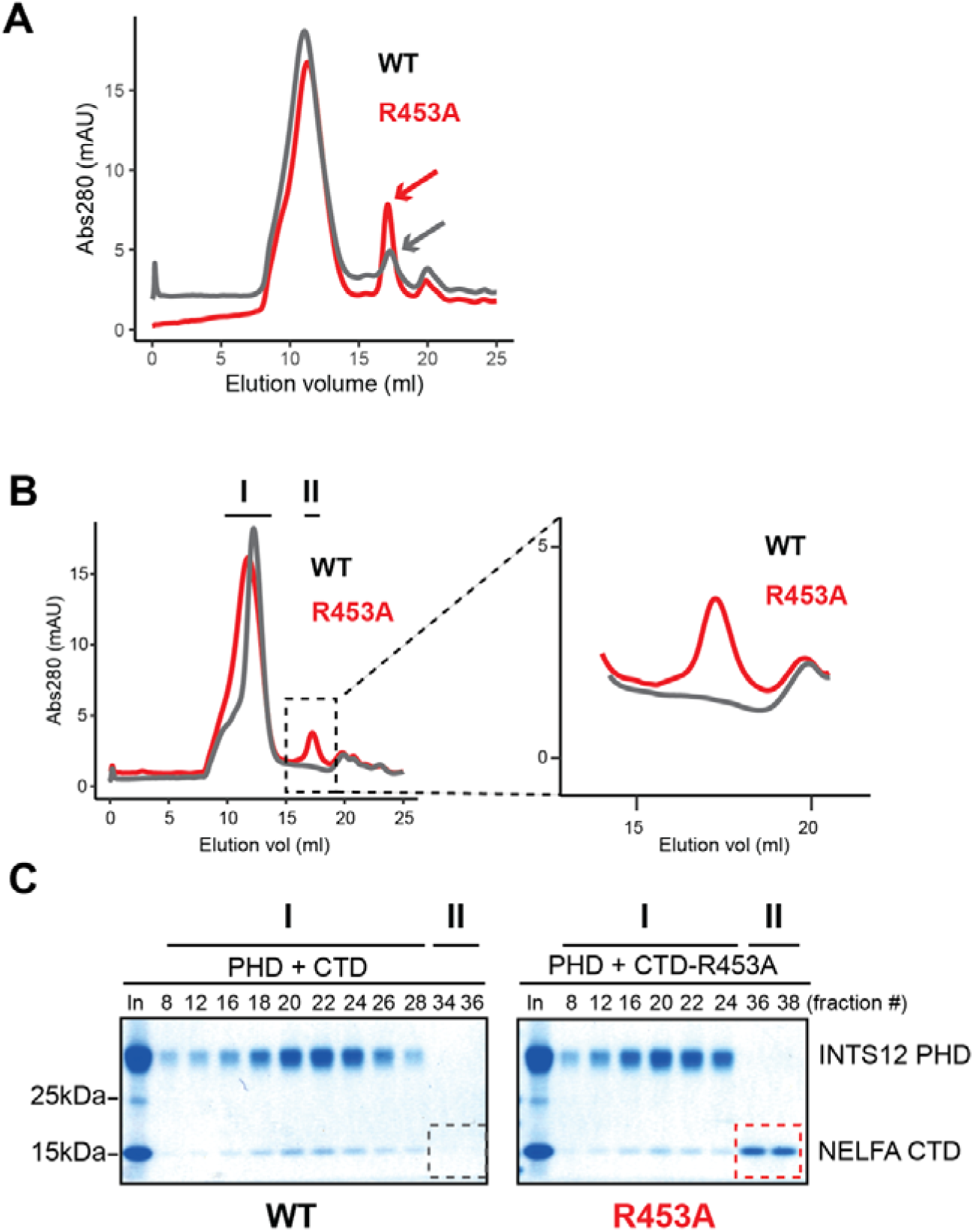
**A.** UV absorption of Superdex 200 for samples in which GST-INTS12 PHD was pre-incubated with either NELFA CTD WT or R453A mutant domain under “high” salt conditions (150 mM NaCl), as in Figure 5E. **B.** Similar to A, but with pre-incubation performed under “low” salt conditions (50 mM) NaCl). **C.** Instant Blue staining of the respective Superdex 200 peak fractions from I and II from B.

**Supplementary Figure S7, related to Figure 5.**
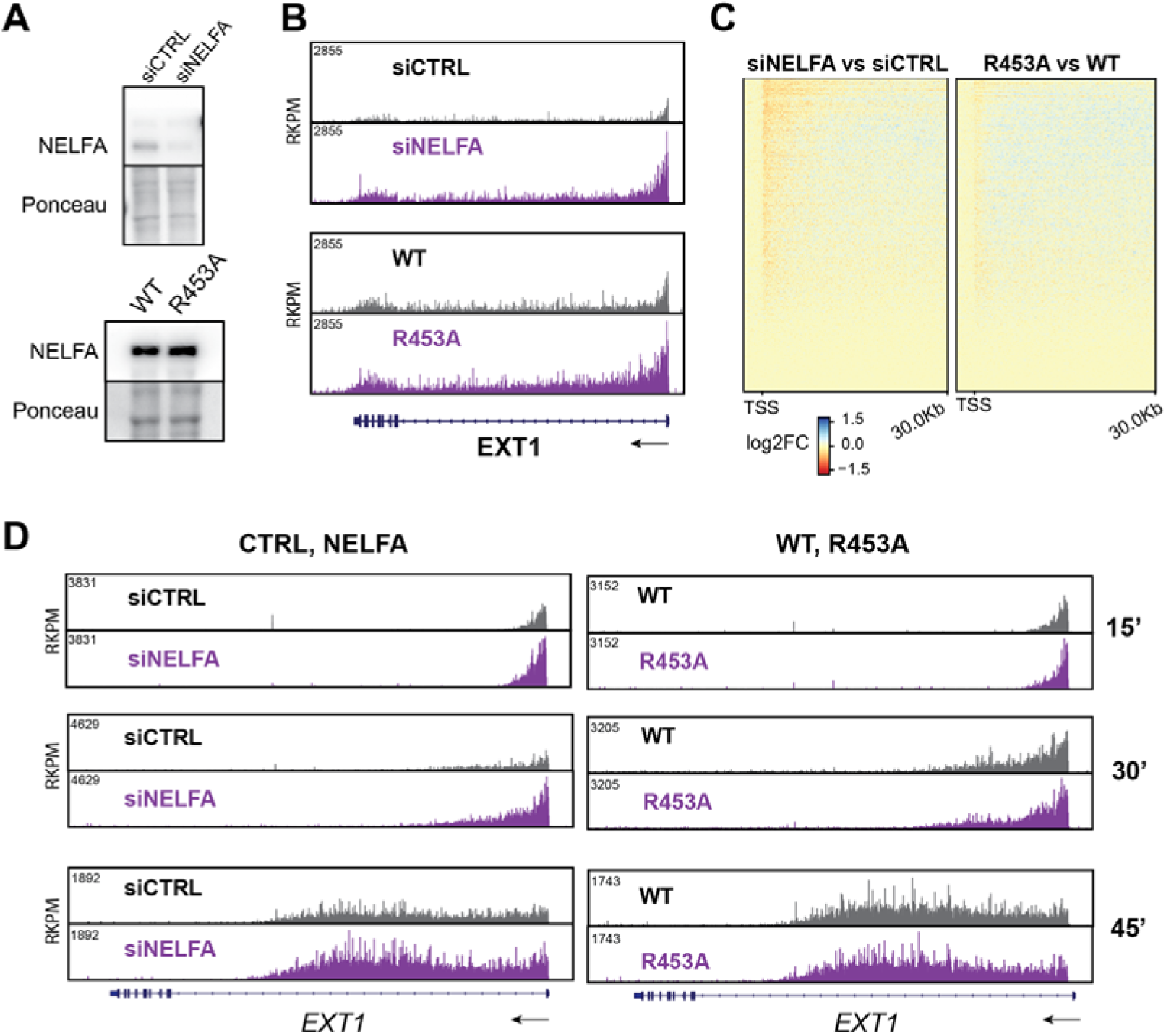
**A.** Western blot of NELFA showing efficiency of knockdown. **B.** Single-gene TTchem-seq example of *EXT1* in NELFA knockdown, as well as in NELFA WT and R453A switchover systems. **C.** Heatmaps showing log2 fold-change in TTchem-seq signal across the first 30kb of protein-coding genes. Log2FC is given for NELFA knockdown vs control, and NELFA R453A vs WT switchovers. Data is aligned at the TSS. **D.** Single gene example of TTchem-seq signal on *EXT1* at several timepoints after DRB washout.

**Supplementary Figure S8, related to Figure 5.**
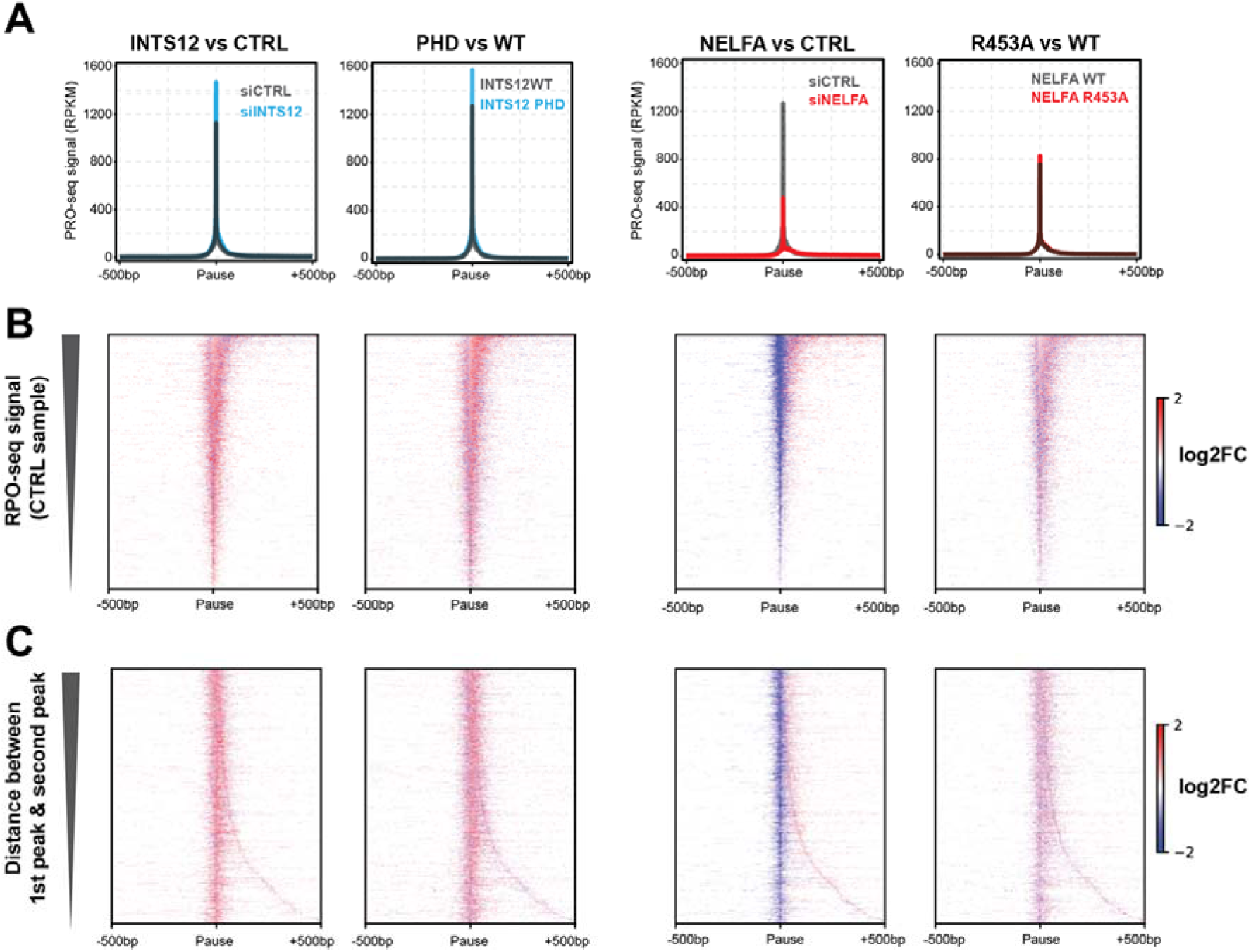
**A.** Metagene profiles aligned on promoter-proximal pause peaks. **B.** Difference maps showing log2FC between respective samples. Genes are ranked according to PRO-seq signal in the Control sample. **C.** Difference maps showing log2FC between respective samples. Genes are ranked by distance between the 1st and second pause positions.

## References

1. Wagner, E.J., Tong, L., and Adelman, K. (2023). Integrator is a global promoter-proximal termination complex. Mol Cell 83, 416–427. 10.1016/j.molcel.2022.11.012.

2. Blears, D., and Svejstrup, J.Q. (2025). Transcription quality control at the promoter-proximal checkpoint. Genes Dev 39, 1399–1413. 10.1101/gad.352973.125.

3. Evans, R., Weber, J., Ziff, E., and Darnell, J.E. (1979). Premature termination during adenovirus transcription. Nature 278, 367–370. 10.1038/278367a0.

4. Marshall, N.F., and Price, D.H. (1992). Control of formation of two distinct classes of RNA polymerase II elongation complexes. Mol Cell Biol 12, 2078–2090.

5. Krebs, A.R., Imanci, D., Hoerner, L., Gaidatzis, D., Burger, L., and Schubeler, D. (2017). Genome-wide Single-Molecule Footprinting Reveals High RNA Polymerase II Turnover at Paused Promoters. Mol Cell 67, 411–422 e414. 10.1016/j.molcel.2017.06.027.

6. Steurer, B., Janssens, R.C., Geverts, B., Geijer, M.E., Wienholz, F., Theil, A.F., Chang, J., Dealy, S., Pothof, J., van Cappellen, W.A., et al. (2018). Live-cell analysis of endogenous GFP-RPB1 uncovers rapid turnover of initiating and promoter-paused RNA Polymerase II. Proc Natl Acad Sci U S A 115, E4368–E4376. 10.1073/pnas.1717920115.

7. Wada, T., Takagi, T., Yamaguchi, Y., Ferdous, A., Imai, T., Hirose, S., Sugimoto, S., Yano, K., Hartzog, G.A., Winston, F., et al. (1998). DSIF, a novel transcription elongation factor that regulates RNA polymerase II processivity, is composed of human Spt4 and Spt5 homologs. Genes Dev 12, 343–356. 10.1101/gad.12.3.343.

8. Yamaguchi, Y., Takagi, T., Wada, T., Yano, K., Furuya, A., Sugimoto, S., Hasegawa, J., and Handa, H. (1999). NELF, a multisubunit complex containing RD, cooperates with DSIF to repress RNA polymerase II elongation. Cell 97, 41–51. 10.1016/s0092-8674(00)80713-8.

9. Elrod, N.D., Henriques, T., Huang, K.L., Tatomer, D.C., Wilusz, J.E., Wagner, E.J., and Adelman, K. (2019). The Integrator Complex Attenuates Promoter-Proximal Transcription at Protein-Coding Genes. Mol Cell 76, 738–752 e737. 10.1016/j.molcel.2019.10.034.

10. Tatomer, D.C., Elrod, N.D., Liang, D., Xiao, M.S., Jiang, J.Z., Jonathan, M., Huang, K.L., Wagner, E.J., Cherry, S., and Wilusz, J.E. (2019). The Integrator complex cleaves nascent mRNAs to attenuate transcription. Genes Dev 33, 1525–1538. 10.1101/gad.330167.119.

11. Aoi, Y., Smith, E.R., Shah, A.P., Rendleman, E.J., Marshall, S.A., Woodfin, A.R., Chen, F.X., Shiekhattar, R., and Shilatifard, A. (2020). NELF Regulates a Promoter-Proximal Step Distinct from RNA Pol II Pause-Release. Mol Cell 78, 261–274 e265. 10.1016/j.molcel.2020.02.014.

12. Zheng, H., Qi, Y., Hu, S., Cao, X., Xu, C., Yin, Z., Chen, X., Li, Y., Liu, W., Li, J., et al. (2020). Identification of Integrator-PP2A complex (INTAC), an RNA polymerase II phosphatase. Science 370. 10.1126/science.abb5872.

13. Vervoort, S.J., Welsh, S.A., Devlin, J.R., Barbieri, E., Knight, D.A., Offley, S., Bjelosevic, S., Costacurta, M., Todorovski, I., Kearney, C.J., et al. (2021). The PP2A-Integrator-CDK9 axis fine-tunes transcription and can be targeted therapeutically in cancer. Cell 184, 3143–3162 e3132. 10.1016/j.cell.2021.04.022.

14. Welsh, S.A., and Gardini, A. (2023). Genomic regulation of transcription and RNA processing by the multitasking Integrator complex. Nat Rev Mol Cell Biol 24, 204–220. 10.1038/s41580-022-00534-2.

15. Song, A., Lu, D., and Chen, F.X. (2025). The Integrator-PP2A complex integrates promoter-proximal premature termination with chromatin context and genome maintenance. Trends Biochem Sci 50, 1118–1130. 10.1016/j.tibs.2025.09.006.

16. Vos, S.M., Farnung, L., Urlaub, H., and Cramer, P. (2018). Structure of paused transcription complex Pol II-DSIF-NELF. Nature 560, 601–606. 10.1038/s41586-018-0442-2.

17. Fianu, I., Chen, Y., Dienemann, C., Dybkov, O., Linden, A., Urlaub, H., and Cramer, P. (2021). Structural basis of Integrator-mediated transcription regulation. Science 374, 883–887. 10.1126/science.abk0154.

18. Fianu, I., Ochmann, M., Walshe, J.L., Dybkov, O., Cruz, J.N., Urlaub, H., and Cramer, P. (2024). Structural basis of Integrator-dependent RNA polymerase II termination. Nature 629, 219–227. 10.1038/s41586-024-07269-4.

19. Li, H., Ilin, S., Wang, W., Duncan, E.M., Wysocka, J., Allis, C.D., and Patel, D.J. (2006). Molecular basis for site-specific read-out of histone H3K4me3 by the BPTF PHD finger of NURF. Nature 442, 91–95. nature04802 [pii] 10.1038/nature04802.

20. Shi, X., Hong, T., Walter, K.L., Ewalt, M., Michishita, E., Hung, T., Carney, D., Pena, P., Lan, F., Kaadige, M.R., et al. (2006). ING2 PHD domain links histone H3 lysine 4 methylation to active gene repression. Nature 442, 96–99. nature04835 [pii] 10.1038/nature04835.

21. Wysocka, J., Swigut, T., Xiao, H., Milne, T.A., Kwon, S.Y., Landry, J., Kauer, M., Tackett, A.J., Chait, B.T., Badenhorst, P., et al. (2006). A PHD finger of NURF couples histone H3 lysine 4 trimethylation with chromatin remodelling. Nature 442, 86–90. nature04815 [pii] 10.1038/nature04815.

22. Vermeulen, M., Mulder, K.W., Denissov, S., Pijnappel, W.W., van Schaik, F.M., Varier, R.A., Baltissen, M.P., Stunnenberg, H.G., Mann, M., and Timmers, H.T. (2007). Selective anchoring of TFIID to nucleosomes by trimethylation of histone H3 lysine 4. Cell 131, 58–69. S0092-8674(07)01079-3 [pii] 10.1016/j.cell.2007.08.016.

23. Sanchez, R., and Zhou, M.M. (2011). The PHD finger: a versatile epigenome reader. Trends Biochem Sci 36, 364–372. 10.1016/j.tibs.2011.03.005.

24. Jain, K., Fraser, C.S., Marunde, M.R., Parker, M.M., Sagum, C., Burg, J.M., Hall, N., Popova, I.K., Rodriguez, K.L., Vaidya, A., et al. (2020). Characterization of the plant homeodomain (PHD) reader family for their histone tail interactions. Epigenetics Chromatin 13, 3. 10.1186/s13072-020-0328-z.

25. Blears, D., Lou, J., Fong, N., Mitter, R., Sheridan, R.M., He, D., Dirac-Svejstrup, A.B., Bentley, D., and Svejstrup, J.Q. (2024). Redundant pathways for removal of defective RNA polymerase II complexes at a promoter-proximal pause checkpoint. Mol Cell 84, 4790–4807 e4711. 10.1016/j.molcel.2024.10.012.

26. Fong, N., Sheridan, R.M., Ramachandran, S., and Bentley, D.L. (2022). The pausing zone and control of RNA polymerase II elongation by Spt5: Implications for the pause-release model. Mol Cell 82, 3632–3645 e3634. 10.1016/j.molcel.2022.09.001.

27. Stein, C.B., Field, A.R., Mimoso, C.A., Zhao, C., Huang, K.L., Wagner, E.J., and Adelman, K. (2022). Integrator endonuclease drives promoter-proximal termination at all RNA polymerase II-transcribed loci. Mol Cell 82, 4232–4245 e4211. 10.1016/j.molcel.2022.10.004.

28. Hu, S., Peng, L., Song, A., Ji, Y.X., Cheng, J., Wang, M., and Chen, F.X. (2023). INTAC endonuclease and phosphatase modules differentially regulate transcription by RNA polymerase II. Mol Cell 83, 1588–1604 e1585. 10.1016/j.molcel.2023.03.022.

29. Gregersen, L.H., Mitter, R., and Svejstrup, J.Q. (2020). Using TT(chem)-seq for profiling nascent transcription and measuring transcript elongation. Nat Protoc 15, 604–627. 10.1038/s41596-019-0262-3.

30. Lykke-Andersen, S., Zumer, K., Molska, E.S., Rouviere, J.O., Wu, G., Demel, C., Schwalb, B., Schmid, M., Cramer, P., and Jensen, T.H. (2021). Integrator is a genome-wide attenuator of non-productive transcription. Mol Cell 81, 514–529 e516. 10.1016/j.molcel.2020.12.014.

31. Baluapuri, A., Zhao, N.C., Marina, R.J., Huang, K.L., Kuzkina, A., Amodeo, M.E., Stein, C.B., Ahn, L.Y., Farr, J.S., Schaffer, A.E., et al. (2025). Integrator loss leads to dsRNA formation that triggers the integrated stress response. Cell 188, 3184–3201.e3121. 10.1016/j.cell.2025.03.025.

32. Baillat, D., Hakimi, M.A., Naar, A.M., Shilatifard, A., Cooch, N., and Shiekhattar, R. (2005). Integrator, a multiprotein mediator of small nuclear RNA processing, associates with the C-terminal repeat of RNA polymerase II. Cell 123, 265–276. S0092-8674(05)00824-X [pii] 10.1016/j.cell.2005.08.019.

33. Ezzeddine, N., Chen, J., Waltenspiel, B., Burch, B., Albrecht, T., Zhuo, M., Warren, W.D., Marzluff, W.F., and Wagner, E.J. (2011). A subset of Drosophila integrator proteins is essential for efficient U7 snRNA and spliceosomal snRNA 3’-end formation. Mol Cell Biol 31, 328–341. 10.1128/MCB.00943-10.

34. Chen, J., Waltenspiel, B., Warren, W.D., and Wagner, E.J. (2013). Functional Analysis of the Integrator Subunit 12 Identifies a Microdomain That Mediates Activation of the Drosophila Integrator Complex. Journal of Biological Chemistry 288, 4867–4877. 10.1074/jbc.M112.425892.

35. Stadelmayer, B., Micas, G., Gamot, A., Martin, P., Malirat, N., Koval, S., Raffel, R., Sobhian, B., Severac, D., Rialle, S., et al. (2014). Integrator complex regulates NELF-mediated RNA polymerase II pause/release and processivity at coding genes. Nat Commun 5, 5531. 10.1038/ncomms6531.

36. Yamamoto, J., Hagiwara, Y., Chiba, K., Isobe, T., Narita, T., Handa, H., and Yamaguchi, Y. (2014). DSIF and NELF interact with Integrator to specify the correct post-transcriptional fate of snRNA genes. Nat Commun 5, 4263. 10.1038/ncomms5263.

37. Aoi, Y., Smith, E.R., Shah, A.P., Rendleman, E.J., Marshall, S.A., Woodfin, A.R., Chen, F.X., Shiekhattar, R., and Shilatifard, A. (2020). NELF Regulates a Promoter-Proximal Step Distinct from RNA Pol II Pause-Release. Mol Cell 78, 261–274. 10.1016/j.molcel.2020.02.014.

38. Judd, J., Wojenski, L.A., Wainman, L.M., Tippens, N.D., Rice, E.J., Dziubek, A., Villafano, G.J., Wissink, E.M., Versluis, P., Bagepalli, L., et al. (2020). A rapid, sensitive, scalable method for Precision Run-On sequencing (PRO-seq). bioRxiv 2020.05.18.102277; doi: 10.1101/2020.05.18.102277.

39. Aoi, Y., Takahashi, Y.H., Shah, A.P., Iwanaszko, M., Rendleman, E.J., Khan, N.H., Cho, B.K., Goo, Y.A., Ganesan, S., Kelleher, N.L., and Shilatifard, A. (2021). SPT5 stabilization of promoter-proximal RNA polymerase II. Mol Cell 81, 4413–4424 e4415. 10.1016/j.molcel.2021.08.006.

40. Gaurav, N., and Kutateladze, T.G. (2023). Non-histone binding functions of PHD fingers. Trends Biochem Sci 48, 610–617. 10.1016/j.tibs.2023.03.005.

41. Saibil, H.R. (2022). Cryo-EM in molecular and cellular biology. Mol Cell 82, 274–284. 10.1016/j.molcel.2021.12.016.

42. Cermakova, K., and Hodges, H.C. (2023). Interaction modules that impart specificity to disordered protein. Trends Biochem Sci 48, 477–490. 10.1016/j.tibs.2023.01.004.

43. Meier, F., Brunner, A.D., Koch, S., Koch, H., Lubeck, M., Krause, M., Goedecke, N., Decker, J., Kosinski, T., Park, M.A., et al. (2018). Online Parallel Accumulation-Serial Fragmentation (PASEF) with a Novel Trapped Ion Mobility Mass Spectrometer. Mol Cell Proteomics 17, 2534–2545. 10.1074/mcp.TIR118.000900.

44. Nojima, T., Gomes, T., Grosso, A.R., Kimura, H., Dye, M.J., Dhir, S., Carmo-Fonseca, M., and Proudfoot, N.J. (2015). Mammalian NET-Seq Reveals Genome-wide Nascent Transcription Coupled to RNA Processing. Cell 161, 526–540. 10.1016/j.cell.2015.03.027.

45. Li, B., and Dewey, C.N. (2011). RSEM: accurate transcript quantification from RNA-Seq data with or without a reference genome. BMC Bioinformatics 12, 323. 10.1186/1471-2105-12-323.

46. Dobin, A., Davis, C.A., Schlesinger, F., Drenkow, J., Zaleski, C., Jha, S., Batut, P., Chaisson, M., and Gingeras, T.R. (2013). STAR: ultrafast universal RNA-seq aligner. Bioinformatics 29, 15–21. 10.1093/bioinformatics/bts635.

47. Mirdita, M., Schutze, K., Moriwaki, Y., Heo, L., Ovchinnikov, S., and Steinegger, M. (2022). ColabFold: making protein folding accessible to all. Nat Methods 19, 679–682. 10.1038/s41592-022-01488-1.

